# Trial history and model-derived reward prediction errors in mouse visual decisions

**DOI:** 10.64898/2026.11.19.767881

**Authors:** Amitis Mirabedini, Raya JanatAbadi, Alireza Khanteymoori

**Affiliations:** Electrical Engineering Department, Sharif University of Technology, Tehran, Iran; Computer Engineering Department, Sharif University of Technology, Tehran, Iran; Department of Psychology, University of Freiburg, Freiburg, Germany

## Abstract

A reward prediction error (RPE) estimated from an animal’s behavior can predict its next choice. Part of this predictive value may come from information about earlier trials that a choice model could also use directly. We asked whether a model-derived RPE improves choice prediction beyond the observable history of stimuli, choices, and rewards. We analyzed 275,292 trials from 107 mice performing the International Brain Laboratory visual decision task and evaluated the main choice models on sessions that were not used for fitting. Recent choices strongly improved prediction. An RPE based on a stimulus-dependent reward expectation added a small gain beyond the preceding choice and reward, and this gain fell to near zero once the model included the preceding stimulus. When the model also summarized choices and rewards over several earlier trials, an RPE trace with the same retention parameter as the history traces did not reliably improve prediction. In simulated agents with known decision rules, a simplified version of this comparison detected an RPE-based learning rule in which choices depended on accumulated prediction errors, and did not improve prediction for an agent that used trial history directly. In mice, the state-dependent RPE and its trace added little predictive information beyond observable trial history, which limits what choice prediction alone can reveal about prediction-error learning.

## Introduction

The same reward can be expected in one context and surprising in another. In reinforcement learning, a reward prediction error (RPE) is the difference between received and expected reward. Phasic activity in midbrain dopamine neurons has been linked to this discrepancy [1, 2]. We asked whether an RPE estimated from behavior improves choice prediction after the model accounts for recent trials. In perceptual decisions, sensory evidence can shape how confidently an animal expects a choice to be rewarded. Reward has been shown to bias subsequent perceptual choices particularly after difficult decisions [3].

This dependence on perceptual context complicates the interpretation of a model-derived RPE. For example, a model that assigns a lower probability of reward to a choice made under weak sensory evidence will assign a larger positive prediction error when that choice is rewarded. The resulting RPE therefore carries information about the previous stimulus as well as the outcome. In the task we analyzed, the reward is determined by the stimulus side and the choice whenever the contrast is nonzero, so the previous choice and reward already indicate on which side the previous stimulus appeared. What a model of choice and reward history lacks is mainly the strength of the previous stimulus, that is, how difficult the previous decision was. If a model gains predictive performance after adding this RPE, some of that gain may come from information about the previous stimulus that was absent from the reference model. Including the preceding stimulus directly tests whether the RPE adds predictive value after the model has access to that information.

### Related work

Previous choices and outcomes can influence perceptual decisions in ways that depend on the structure of the task. In two-choice tasks with sequential regularities, rats relied more strongly on accumulated evidence about the stimulus sequence after correct responses than after errors [4, 5]. In the International Brain Laboratory’s visual decision-making task, an exponentially weighted history of recent actions better accounted for mice’s choices than models based on recent stimuli or a Bayes-optimal estimate of the stimulus prior [6]. Together,these findings motivate a reference model in which past choices can influence the current decision and that influence can depend on whether those choices were rewarded.

The comparison also depends on how far back each predictor reaches. Fritsche et al. showed that a reinforcement-learning model with multi-trial memory could account for how past choices influenced current choices [7]. Their findings motivate asking how much a modeled RPE adds when the reference model can already use earlier trials. A reward prediction error computed from learned values can likewise carry information from trials earlier than the one on which it is computed, because the action values of a reinforcement-learning model are determined by the history of past choices and rewards [8]. In mice performing a probabilistic two-armed bandit task, a forgetting Q-learning model and a recursively formulated logistic-regression model were shown to yield mathematically equivalent trial-by-trial choice probabilities [9]. An apparent predictive advantage of an RPE-derived predictor over a one-trial history model could therefore arise from information carried across earlier trials.

### Overview of the analyses

We addressed this question in the International Brain Laboratory (IBL) visual decision task [10] through a sequence of analyses (Fig 1C). We first screened six candidate signals across all 139 mice in the dataset. Five were derived from a Q-learning model, and one, the change in outcome between consecutive trials, required no learning model. This screen fitted and compared the models on the same trials and served only to describe the candidates.

**Fig 1.**
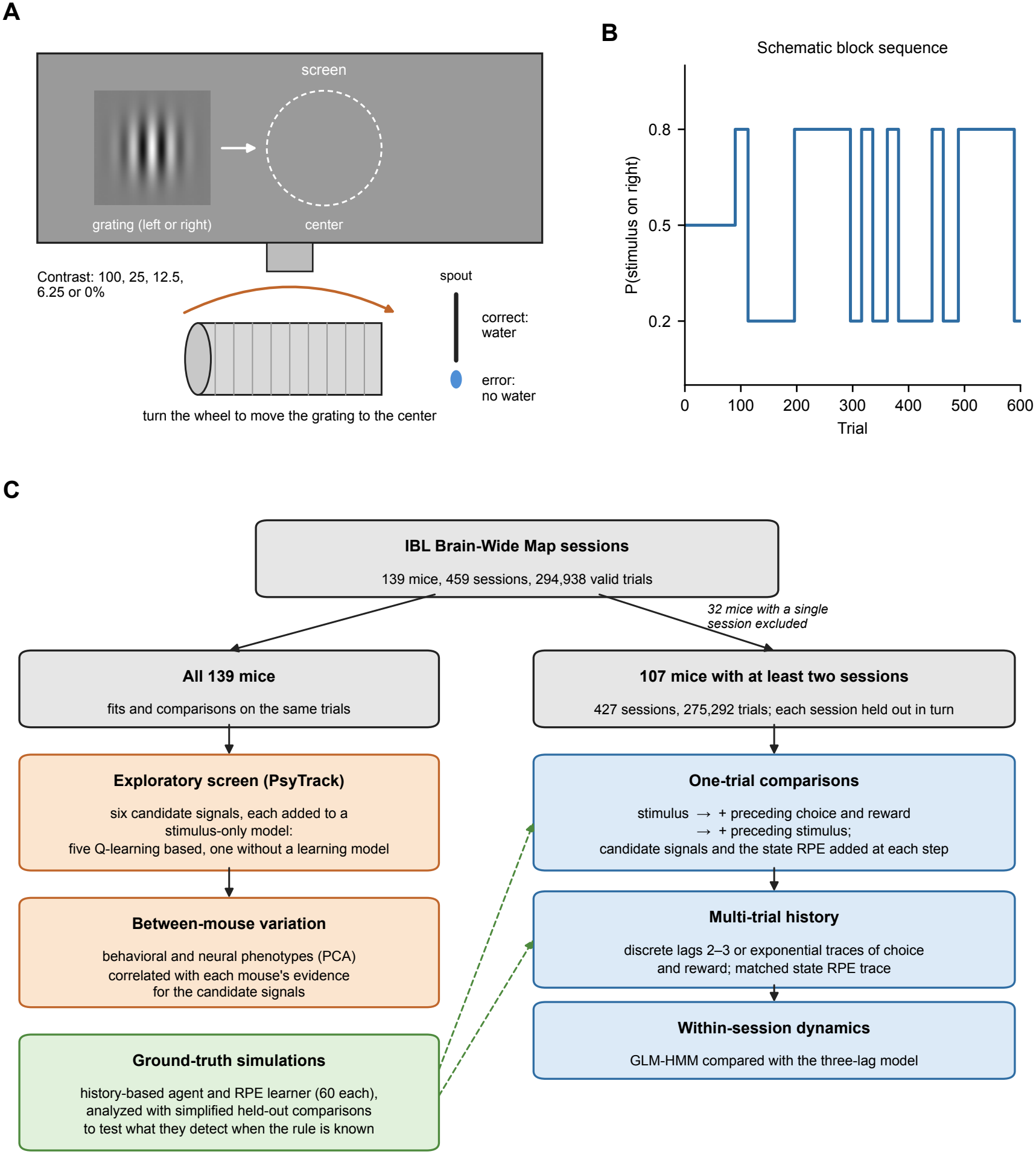
Task and overview of the analyses. (A) Schematic of the IBL visual decision task. A grating appeared on the left or right of a screen, and the head-fixed mouse turned a wheel to move it to the center. Correct choices were rewarded with water. Grating contrast varied across trials. (B) Schematic sequence of the programmed probability that the stimulus appears on the right. Each session began with an unbiased block (probability 0.5), followed by unsignaled blocks in which the probability alternated between 0.2 and 0.8. The block lengths shown are illustrative.(C) Analyses in this study. Exploratory analyses (orange) used all 139 mice and fitted and compared models on the same trials. The main analyses (blue) used the 107 mice with at least two sessions and evaluated each model on held-out sessions. Simulated agents with known decision rules (green) were analyzed with simplified versions of the held-out comparisons.

The main analyses evaluated choice prediction in held-out sessions from the 107 mice with at least two sessions. Starting from a model with the current stimulus, we added the preceding choice and reward and then the preceding stimulus, and at each step tested whether an RPE-derived predictor improved prediction. For this purpose, we defined a state-dependent RPE with a reward expectation conditioned on the stimulus and chosen action. This formulation allowed us to test whether its predictive contribution reflected information about the preceding stimulus. We then extended the reference models to several earlier trials, using discrete lags or exponentially weighted traces, and compared an RPE trace with choice and reward traces of the same retention.

Two further analyses examined the structure of choice behavior. A generalized linear model with hidden Markov states (GLM-HMM) tested whether allowing the decision strategy to change within a session improved prediction beyond a fixed multi-trial model. A between-mouse analysis tested whether evidence for the candidate signals covaried with behavioral and neural phenotypes. Finally, we simulated agents with known decision rules and analyzed them with simplified versions of the same comparisons, to determine what these comparisons can detect when the mechanism is known. The state-dependent RPE added little predictive information once the reference models included the preceding stimulus and longer choice and reward histories.

## Materials and methods

### Behavioral task

We analyzed choices from mice performing the International Brain Laboratory’s visual decision-making task [10]. On each trial, a grating appeared to the left or right of the center of a screen. Head-fixed mice turned a wheel to move the grating toward the center, thereby reporting its location. Correct choices were rewarded with water. Grating contrast varied across trials, providing different strengths of sensory evidence about the stimulus location. The probability of a right-side stimulus was 0.5 in unbiased blocks and either 0.2 or 0.8 in biased blocks. Block transitions were not explicitly signaled to the mouse (Fig 1A and 1B).

We chose this task because the stimulus, choice, and reward are recorded on each trial. We could therefore test whether a model-derived RPE helps predict the next choice after the reference model already includes the preceding stimulus, choice, and reward. The recorded stimulus-side probabilities also allowed us to test whether any gain persisted after accounting for the task’s block structure. Repeated sessions from the same mouse allowed us to fit models on one set of sessions and evaluate them on another. Variation in stimulus contrast also allowed us to test whether the RPE’s predictive contribution reflected the strength of the preceding sensory evidence.

### Dataset construction

We constructed the trial and phenotype datasets from sessions in the IBL Brain-Wide Map [11]. Table 1 lists every field we used, its source, and the variable we derived from it.

**Table 1.** Fields taken from the IBL Brain-Wide Map data and the variables derived from them.

| Source | Field | Derived variable and use |
|---|---|---|
| *A. Trial dataset (DANDI 000409, NWB trials table)* |  |  |
| NWB trials table | mouse_wheel_choice | Choice *c<sub>t</sub>* (+1 right, −1 left); trials without a response excluded |
|  | gabor_stimulus_side,<br>gabor_stimulus_contrast | Left and right contrasts, transformed with Eq (2); signed stimulus *s̃<sub>t</sub>* |
|  | is_mouse_rewarded | Reward *r<sub>t</sub>* (1 or 0) and centered reward 2*r<sub>t</sub>* − 1 |
|  | probability_left | Unbiased, left-biased, and right-biased block indicators (block-control analysis) |
| NWB file | session_start_time | Chronological order of the sessions of each mouse |
| *B. Session datasets for the phenotypes (IBL public data server, same 459 sessions)* |  |  |
| Trials table | choice, contrasts, feedback type, block probability, stimulus-onset, go-cue, first-movement, response, and feedback times | Performance, psychometric, history, block-prior, and timing features |
| Wheel | wheel position and time stamps; wheel-movement intervals and peak amplitudes | Wheel features |
| Left and right cameras | pupil diameter (smoothed) and frame times | Pupil features |
|  | region-of-interest motion energy | Motion-energy features |
| Spike sorting (revision 2024-05-06) | cluster quality label and firing rate, spike amplitudes, waveform peak-to-trough duration | Neural features of units passing all quality metrics |
|  | cluster channels and channel brain locations | Assignment of units to Beryl atlas regions |

The trial dataset contains 294,938 trials from 459 sessions and 139 mice. The derived trial-level variables, the mouse-level behavioral and neural features, and the per-mouse results are listed in the Data availability statement.

#### Trial dataset for the choice models

We took the trial tables from the processed behavior and electro-physiology Neurodata Without Borders (NWB) files of DANDI Dandiset 000409, version 0.260309.1324 [12], one file per session. This version contained 459 such sessions from 139 mice. We streamed each file with the DANDI and PyNWB Python packages and read only its trials table. From it we derived the choice (*c*_*t*_ = +1 for a rightward and *−*1 for a leftward choice), the left and right contrasts (the recorded contrast on the stimulus side and zero on the other side), and the reward (*r*_*t*_ = 1 for rewarded and 0 for unrewarded trials). The recorded block probability was used only in the block-control analysis. As a consistency check, on every trial with a nonzero contrast the recorded reward matched whether the derived choice was on the stimulus side.

We retained trials with a left or right response, interpretable feedback, and nonmissing contrast values. Of the 296,090 recorded trials, 1,152 (0.39%) were excluded, all of them trials without a response. We required at least 10 valid trials per session and at least 50 valid trials per mouse, and all 459 sessions met these criteria. The sessions of each mouse were ordered by their recorded start time and concatenated, and all history predictors and learning models were reset at session boundaries. We constructed history predictors after filtering, so the previous trial refers to the preceding retained trial in the same session. Because only trials without a response were removed, the preceding retained trial differed from the preceding recorded trial only after these 1,152 trials.

The resulting dataset comprised 294,938 valid trials from 459 sessions across 139 mice. The exploratory PsyTrack and mouse-level phenotype analyses used all 139 mice. The held-out choice analyses required at least two usable sessions per mouse, because each session was predicted by a model fitted to the other sessions of the same mouse. Thirty-two mice contributed a single session (S7 Fig), which left 275,292 trials from 427 sessions across 107 mice for the held-out analyses.

#### Session datasets for the phenotypes

For the behavioral and neural phenotypes, we used the IBL data files of the same 459 sessions, which belong to the Brain-Wide Map project, through the IBL public data server. These files provide the trial event times, wheel movements, video-derived pupil diameter and motion energy, and the spike-sorting outputs that the NWB trial tables do not contain (Table 1). We used the spike-sorting outputs of the 2024-05-06 revision for all 699 probe insertions.

### Candidate signals

To test whether a reward discrepancy adds predictive value beyond observable history, we constructed candidate signals from the same behavioral records. Each signal compares the outcome of a trial with a reference quantity, and the signal from the preceding trial was used to predict the current choice. We grouped the seven signals by whether they were derived from a reinforcement-learning model (Table 2). Q-learning is a standard model of trial-by-trial value learning [13], and temporal-difference prediction errors have been linked to phasic dopamine responses [1, 2]. Five signals were derived from Q-learning and differed in their learning rule, sign, magnitude, or reference quantity. Two signals did not require a learning model: the change in outcome between consecutive trials and a state-dependent prediction error, whose reward expectation depends on the stimulus and choice. The state-dependent error is the basis of the main held-out analyses.

**Table 2.**
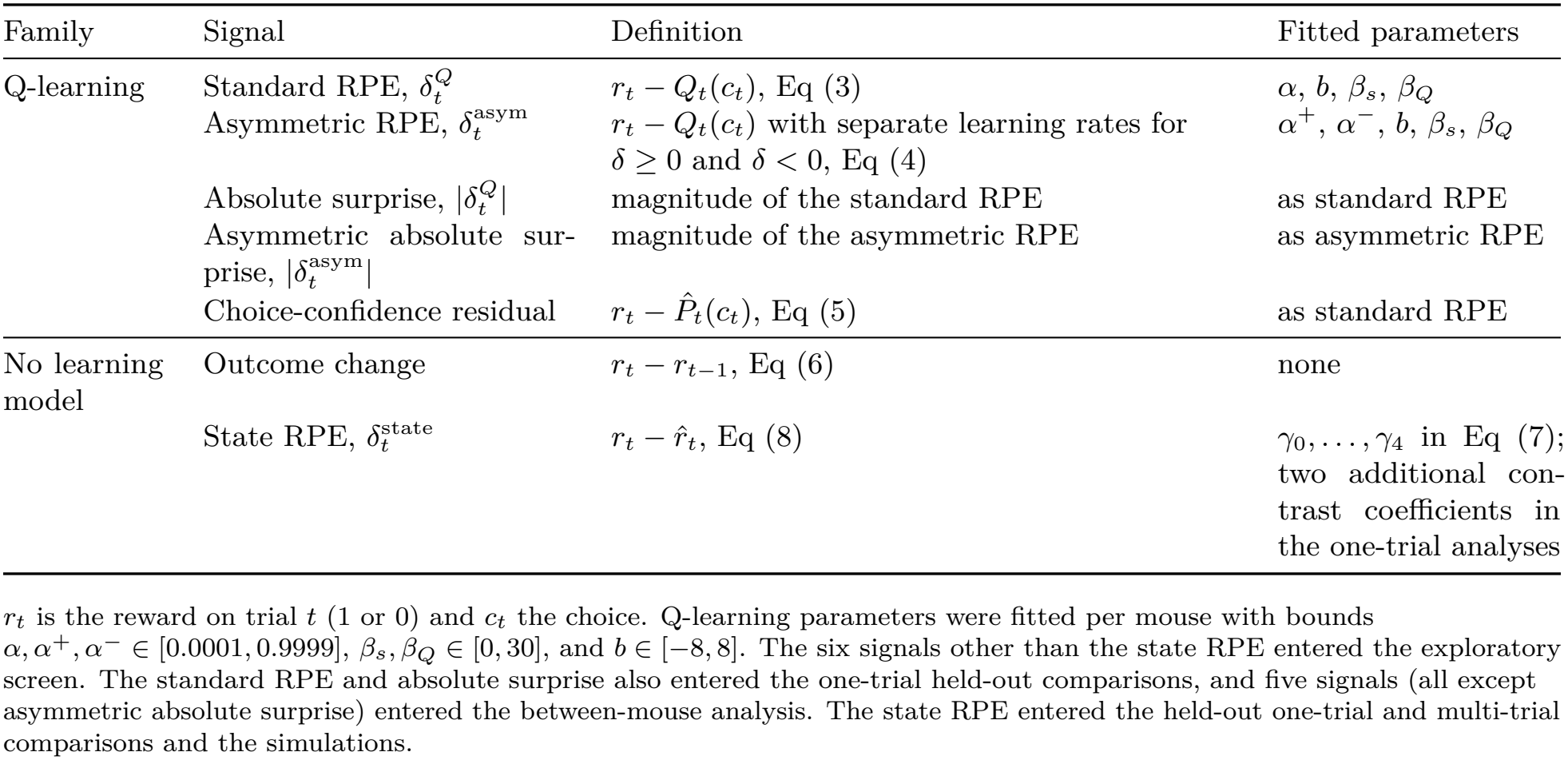
Candidate signals and their fitted parameters.

| Family | Signal | Definition | Fitted parameters |
|---|---|---|---|
| Q-learning | Standard RPE, δ<sub>t</sub><sup>Q</sup> | r<sub>t</sub> − Q<sub>t</sub>(c<sub>t</sub>), Eq (3) | α, b, β<sub>s</sub>, β<sub>Q</sub> |
|  | Asymmetric RPE, δ<sub>t</sub><sup>asym</sup> | r<sub>t</sub> − Q<sub>t</sub>(c<sub>t</sub>) with separate learning rates for δ ≥ 0 and δ < 0, Eq (4) | α<sup>+</sup>, α<sup>−</sup>, b, β<sub>s</sub>, β<sub>Q</sub> |
|  | Absolute surprise, \|δ<sub>t</sub><sup>Q</sup>\| | magnitude of the standard RPE | as standard RPE |
|  | Asymmetric absolute sur-prise, \|δ<sub>t</sub><sup>asym</sup>\| | magnitude of the asymmetric RPE | as asymmetric RPE |
|  | Choice-confidence residual | r<sub>t</sub> − P̂<sub>t</sub>(c<sub>t</sub>), Eq (5) | as standard RPE |
| No learning model | Outcome change | r<sub>t</sub> − r<sub>t−1</sub>, Eq (6) | none |
|  | State RPE, δ<sub>t</sub><sup>state</sup> | r<sub>t</sub> − r̂<sub>t</sub>, Eq (8) | γ<sub>0</sub>, . . . , γ<sub>4</sub> in Eq (7); two additional con-trast coefficients in the one-trial analyses |

r<sub>t</sub> is the reward on trial t (1 or 0) and c<sub>t</sub> the choice. Q-learning parameters were fitted per mouse with bounds α, α<sup>+</sup>, α<sup>−</sup> ∈ [0.0001, 0.9999], β<sub>s</sub>, β<sub>Q</sub> ∈ [0, 30], and b ∈ [−8, 8]. The six signals other than the state RPE entered the exploratory screen. The standard RPE and absolute surprise also entered the one-trial held-out comparisons, and five signals (all except asymmetric absolute surprise) entered the between-mouse analysis. The state RPE entered the held-out one-trial and multi-trial comparisons and the simulations.

#### Signals derived from Q-learning

We used a Q-learning model to estimate the value of each action from the mouse’s previous choices and rewards. Let *Q*_*t*_(*a*) be the value of action *a* before the choice on trial *t*, and let *c*_*t*_ = +1 denote a rightward choice. The model assigned a probability of choosing right given by

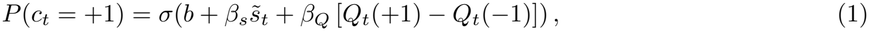

where *σ* is the logistic function, *b* is the choice bias, and *β*_*s*_ and *β*_*Q*_ are the stimulus and action-value weights.

The signed stimulus 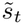 was the transformed right contrast minus the transformed left contrast, using

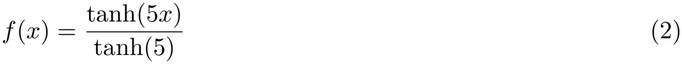

for each contrast. After observing the reward, the model updated the value of the chosen action using the signed prediction error,

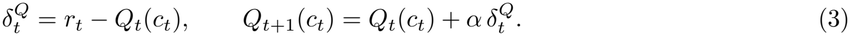

Here, *α* is the learning rate. The unchosen action retained its value, and both action values were initialized to 0.5 at the start of each session.

Separate learning rates for positive and negative prediction errors are a common extension of this model, although an apparent asymmetry can also arise from choice perseveration [14]. The asymmetric model updated the chosen value as

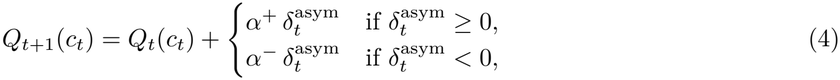

which yielded a second signed signal,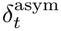 To test whether the magnitude of the discrepancy was predictive regardless of its sign, we also used the absolute values of the two signed errors, 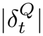and 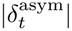, which we refer to as absolute surprise. Unsigned prediction errors of this kind control learning in attention-based learning models [15].

The choice-confidence residual compared the outcome with the probability that the fitted standard Q-learning choice rule, Eq (1), assigned to the action the mouse actually selected,

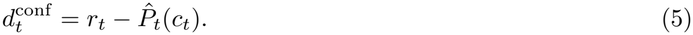

This probability depends on the current stimulus and learned action values. The residual uses the fitted probability of the chosen action as an operational proxy for confidence.

We fitted the learning rates and choice-rule parameters separately for each mouse by maximizing the likelihood of the observed choices with bounded L-BFGS-B from three starting points (bounds in Table 2). For the exploratory screen, the fits used all retained sessions for each mouse.

### Signals without a learning model

The outcome-change signal compared consecutive outcomes directly and required no fitted expectation,

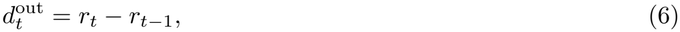

with the preceding reward initialized to 0.5 at the start of each session.

The state-dependent prediction error used a reward expectation conditioned on the stimulus and the chosen action, analogous to the confidence-dependent prediction errors proposed for perceptual decisions [3, 16]. We estimated the probability of reward by logistic regression,

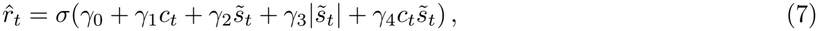

using choice, signed stimulus strength, absolute stimulus strength, and the interaction between choice and signed stimulus strength. In the one-trial analyses, the reward model also included the separate transformed left and right contrasts. Because only one side has a nonzero contrast on each trial, these two terms are linear combinations of the signed and absolute stimulus strength, so they can change the fitted expectation only through regularization. The reward model used standardized predictors and L2 regularization with fixed *C* = 1. Estimated reward probabilities were clipped to [0.001, 0.999] before computing the state-dependent prediction error. The state-dependent prediction error was

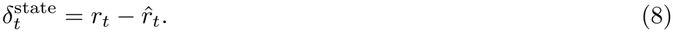

The reward model is fitted to recorded outcomes and does not learn from trial to trial. Its expectation is therefore a model-estimated quantity that depends on the stimulus and choice, not a measurement of the mouse’s own expectation.

### Predicting the next choice

For a candidate signal *d*_*t*_, the corresponding lag-one predictor for the choice on trial *t* was

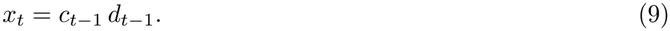

Multiplying by the previous choice expresses the signal relative to the side selected on that trial. We set the lagged predictor to zero on the first trial of each session. In held-out-session analyses, we estimated the parameters of fitted signals using the training data for each split.

For the state-dependent error, this coding separates into three terms. Because the reward is binary, 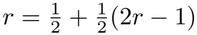, and substituting this into Eq (8) and Eq (9) gives

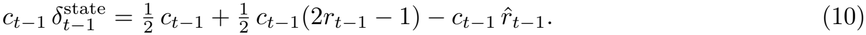

This identity follows from the definitions alone and does not depend on the data. Conceptually, the lagged state RPE combines three pieces of information about the preceding trial: which side was chosen (first term), whether that choice was rewarded (second term), and how much reward the model expected for that choice given the stimulus (third term). The first two terms are already present in the choice and reward history reference model. Once they are included, adding the state-dependent RPE or adding the choice-aligned reward expectation 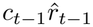 spans the same unpenalized linear predictor space, so the only new information the state RPE can contribute is the reward expectation. With L2 regularization, the two parameterizations can yield different fitted predictions because their coefficients are penalized differently. Because this expectation depends on the preceding stimulus, we compared the state RPE with reference models that included explicit previous-stimulus predictors. These references contained the lagged versions of all inputs to the reward model, Eq (7). Beyond them, the state RPE can therefore add only the nonlinear, choice-aligned function of these inputs that the logistic reward model computes.

### Choice models and held-out comparisons

#### Initial comparison of candidate signals

We used PsyTrack [17] to compare the six candidate signals other than the state RPE across the full cohort while allowing the effects of the current stimulus and the choice bias to vary over trials. PsyTrack models the probability of a rightward choice using logistic regression with trial-specific coefficients,

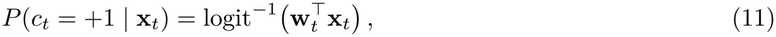

where **x**_*t*_ contains the predictors for trial *t* and **w**_*t*_ contains their coefficients. A Gaussian random-walk prior constrains changes in the coefficients, with separate scales for transitions within and between sessions.

The reference model included a time-varying intercept and separate time-varying coefficients for the transformed left and right stimulus contrasts. We added each lagged, choice-aligned candidate signal separately and compared its model with the reference using the change in approximate log model evidence. Each candidate predictor was z-scored across trials and set to zero on the first trial of each session. We fixed both the within-session and between-session random-walk scales at 2^*−*4^ and the prior scale of the initial weights at 16 across models.

#### Held-out comparisons

We next tested whether the candidate signals improved prediction in held-out sessions. Choice was modeled using L2-regularized logistic regression. The stimulus-only reference included the current transformed left and right contrasts and an intercept. We then added the preceding choice, centered reward, and their interaction to form the one-trial history reference. A further reference included explicit previous-trial stimulus and state variables. At each step, we evaluated an RPE-derived predictor by comparing the model containing it with the corresponding reference on the same held-out trials. This design let us ask how the incremental predictive value of the signal changed as more of its observable inputs entered the reference model. Table 3 lists the predictors and parameters of each reference model.

**Table 3.**
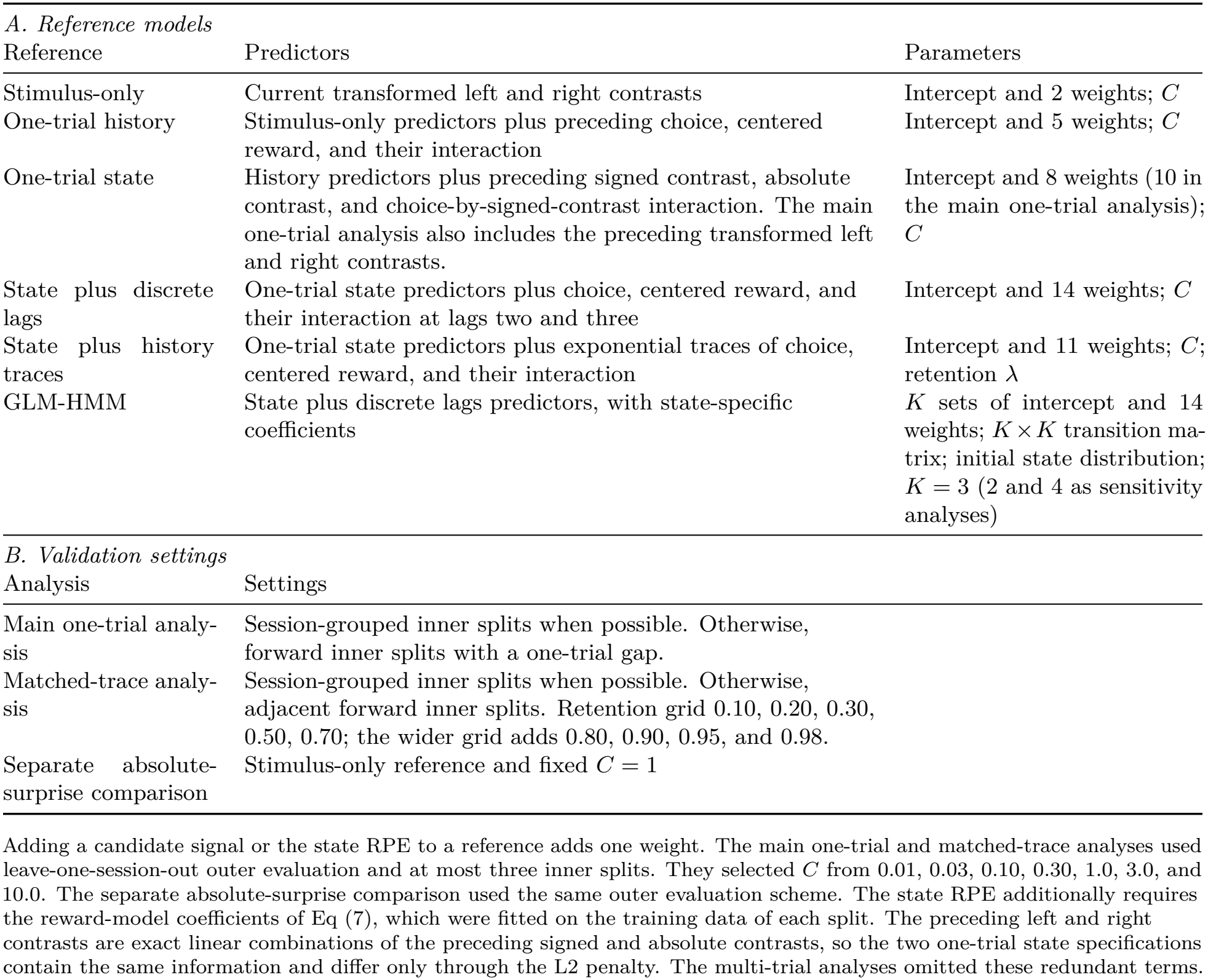
Choice-model reference predictors, parameters, and validation settings.

| *A. Reference models* |  |  |
|---|---|---|
| Reference | Predictors | Parameters |
| Stimulus-only | Current transformed left and right contrasts | Intercept and 2 weights; *C* |
| One-trial history | Stimulus-only predictors plus preceding choice, centered reward, and their interaction | Intercept and 5 weights; *C* |
| One-trial state | History predictors plus preceding signed contrast, absolute contrast, and choice-by-signed-contrast interaction. The main one-trial analysis also includes the preceding transformed left and right contrasts. | Intercept and 8 weights (10 in the main one-trial analysis); *C* |
| State plus discrete lags | One-trial state predictors plus choice, centered reward, and their interaction at lags two and three | Intercept and 14 weights; *C* |
| State plus history traces | One-trial state predictors plus exponential traces of choice, centered reward, and their interaction | Intercept and 11 weights; *C*; retention *λ* |
| GLM-HMM | State plus discrete lags predictors, with state-specific coefficients | *K* sets of intercept and 14 weights; *K* × *K* transition ma-trix; initial state distribution; *K* = 3 (2 and 4 as sensitivity analyses) |
| *B. Validation settings* |  |  |
| Analysis | Settings |  |
| Main one-trial analy-sis | Session-grouped inner splits when possible. Otherwise, forward inner splits with a one-trial gap. |  |
| Matched-trace analy-sis | Session-grouped inner splits when possible. Otherwise, adjacent forward inner splits. Retention grid 0.10, 0.20, 0.30, 0.50, 0.70; the wider grid adds 0.80, 0.90, 0.95, and 0.98. |  |
| Separate absolute-surprise comparison | Stimulus-only reference and fixed *C* = 1 |  |

Adding a candidate signal or the state RPE to a reference adds one weight. The main one-trial and matched-trace analyses used leave-one-session-out outer evaluation and at most three inner splits. They selected *C* from 0.01, 0.03, 0.10, 0.30, 1.0, 3.0, and 10.0. The separate absolute-surprise comparison used the same outer evaluation scheme. The state RPE additionally requires the reward-model coefficients of Eq (7), which were fitted on the training data of each split. The preceding left and right contrasts are exact linear combinations of the preceding signed and absolute contrasts, so the two one-trial state specifications contain the same information and differ only through the L2 penalty. The multi-trial analyses omitted these redundant terms.

For each mouse, we held out one complete session at a time and fitted the models on the remaining sessions. Feature standardization and parameters used to construct fitted predictors were estimated from the training data. In the main one-trial analysis, we refitted the standard Q-learning model within each inner and outer training split. We selected the inverse regularization strength *C* separately for each choice model by maximizing inner-validation log likelihood. We grouped the inner splits by session when multiple training sessions were available. With a single training session, we used forward splits with a one-trial gap. We then refitted the models on all outer-training sessions and scored choices in the held-out session. For Q-learning predictors, earlier choices and rewards in the held-out session updated the action values sequentially, with the fitted parameters fixed.

A separate leave-one-session-out comparison tested absolute surprise against the stimulus-only reference using fixed *C* = 1.

We also tested whether the one-trial results depended on the task’s programmed stimulus prior. We coded left- and right-biased blocks as separate indicators, with the unbiased block as the reference. We added the current block to the stimulus-only model and both the current and preceding block to the history models. We then repeated the matched comparisons on the same held-out sessions, holding the reward model used to construct the state RPE fixed.

### Multi-trial history and matched RPE traces

A state reward prediction error depends on the observed outcome and its expected value. Observable trial history may therefore carry some of the same predictive information as the modeled error. To test how much the state RPE added once the choice model had access to a longer history, we extended the one-trial state reference model in two ways.

Both extensions retained the intercept, the current transformed left and right contrasts, and the one-trial history terms for previous choice, centered reward (2*r*_*t−*1_ *−*1), their interaction, signed and absolute stimulus, and the interaction between choice and signed stimulus. The first extension added choice, centered reward, and their interaction at lags two and three. The second added exponentially weighted traces of the same three variables. For a trial-level variable *x*, the trace available before the choice on trial *t* was

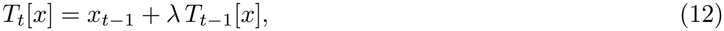

with the trace initialized to zero at the start of each session. The retention parameter *λ* controls how much weight carries forward from earlier trials.

For each reference model, we evaluated two separate additions: the one-trial choice-aligned state RPE and an exponentially weighted trace of the choice-aligned state RPE. The latter applied the same trace definition to 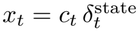. The primary comparison added the state RPE trace to the model with observable-history traces and used the same retention parameter within each fold.

The trace is linear in its input, so the identity in Eq (10) extends to traces with the same retention and zero initialization,

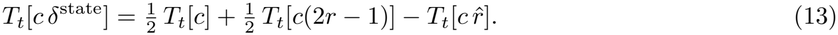

The observable-history reference already includes the first two traces. Adding the state RPE trace or the choice-aligned reward expectation trace therefore gives the same set of unpenalized linear predictors, although regularization can yield different fitted predictions. The reference includes explicit stimulus terms at lag one, whereas the expectation trace can also carry information about stimuli from earlier trials.

We evaluated the models using leave-one-session-out cross-validation. Within each outer training set, we selected *λ* and the logistic regularization parameter *C* jointly using the inner validation log likelihood of the observable-history trace model. We then fixed *λ* and selected *C* separately for each model. Inner validation used session-grouped splits when multiple training sessions were available and adjacent forward splits when only one training session was available. The reward model used to compute the state RPE was fitted on the corresponding training data in each split. Because the original grid often selected its largest value (Results), we repeated the analysis with a wider selection grid. We also evaluated sensitivity to retention using a series of fixed retention values.

### Within-session choice dynamics

The three-lag choice model allowed earlier trials to influence the current choice, with one set of coefficients shared across trials. We tested whether held-out prediction improved when the influence of the stimulus and recent history could vary within a session. We compared the three-lag logistic regression with a GLM-HMM. Both models used the same 14 predictors in the state-plus-discrete-lags reference, plus an intercept (Table 3). History was constructed separately within each session, and unavailable lags at the start of a session were set to zero.

In the GLM-HMM, the latent state *z*_*t*_ determined which set of logistic regression coefficients governed choice on trial *t*. For state *k*, the probability of a rightward choice was

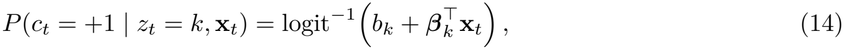

where **x**_*t*_ contains the observed predictors. A first-order Markov chain governed transitions between states,

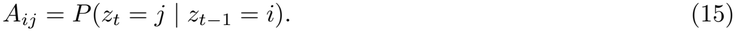

Each session began with the fitted initial state distribution. We used three states for the primary comparison, motivated by prior GLM-HMM work on perceptual choices [18], and evaluated two- and four-state models as sensitivity analyses.

Models were fitted separately for each mouse and evaluated on the same leave-one-session-out folds as the three-lag reference model. This analysis included 107 mice, 427 held-out sessions, and 275,292 trials. In each fold, the GLM-HMM used the predictor scaling and regularization setting selected for the reference model using the training data. We fitted the GLM-HMM by expectation–maximization with two initializations, retaining the fit with the higher training log likelihood. Each fit stopped when the training log likelihood changed by less than 10^*−*6^ per trial or after 35 iterations. After 35 iterations, 258 of 427 three-state fits had not met the convergence criterion. The corresponding counts were 206 for two states and 278 for four states. The held-out gains from these fits could differ from those obtained after convergence.

On each held-out trial, we averaged the state-specific choice probabilities using state probabilities computed from observations through the preceding trial. After scoring the current choice, we used it to update the state probabilities for the next trial. We pooled the held-out log likelihoods across sessions within each mouse and expressed the difference between the GLM-HMM and the reference model per 1,000 trials.

### Behavioral and neural phenotypes

To ask whether differences across mice were associated with the evidence for a candidate signal, we summarized each of the 139 mice with a behavioral phenotype and a neural phenotype. Each mouse contributed one set of phenotype scores and one evidence score for each candidate signal in the association analysis.

#### Behavioral phenotype

We computed mouse-level behavioral features from all sessions of each mouse. Of 35 initial features, six were removed because each nearly duplicated another feature (win-stay, choice entropy, accuracy on hard trials, median response latency, and the pupil and motion responses around feedback). The remaining 29 features covered task performance and psychometric parameters, choice bias and history dependence, sensitivity to the block prior, response and movement timing, wheel activity, pupil responses, and video-derived motion energy (S2 Table). Movement and arousal measures were included because movements and arousal are associated with substantial neural variability and can modulate behavioral performance [19, 20, 21, 22, 23]. Task and wheel features were available for all 139 mice, motion-energy features for 136 mice, and pupil features for 131 mice. We winsorized the features at the 1st and 99th percentiles, imputed missing values with the feature median, and standardized each feature.

Several features depended on recording and protocol factors. Session count, laboratory, and task protocol explained up to 58% of the variance of single features (for example, accuracy), and in an unadjusted principal component analysis (PCA) the first component correlated with the number of sessions (Spearman *ρ* = 0.35, *p* = 2.8 *×*10^*−*5^). We therefore regressed each standardized feature on log session count, laboratory, and task protocol and applied PCA to the residuals. After this adjustment, the correlations of the first two components with session count were near zero (|*ρ*| *<* 0.04). We retained the smallest number of components that together explained at least 80% of the variance.

#### Neural phenotype

Individual differences in behavior have been related to differences in large-scale neural dynamics [24]. We therefore also summarized each mouse by the physiology of the units recorded in its sessions. We used the spike-sorting outputs of the IBL Brain-Wide Map (revision 2024-05-06, Table 1) [11] and retained units that passed all IBL single-unit quality metrics [25]. The 699 probe insertions contained 621,733 units, of which 75,708 were good units. Of these, 65,263 were assigned to a brain region, and the remaining 10,445 were labeled root or void. We assigned units to regions of the Beryl atlas parcellation and retained regions with at least 50 units in total and at least five mice, which left 154 regions containing 94.6% of the anatomically assigned good units.

For each unit we used three quantities: firing rate, extracellular spike amplitude, and waveform peak-to-trough duration. We transformed firing rate as log(1 + *x*), amplitude as log_10_(| *x*| + 10^*−*12^), and peak-to-trough duration as its absolute value. Transformed values were clipped at the 1st and 99th percentiles across anatomically assigned good units. Within each region, values were centered on the median and divided by 1.4826 times the median absolute deviation. For each session–region with at least three units, we calculated the median and interquartile range (IQR). These summaries were combined across sessions within each mouse–region using medians. Each mouse’s relative-level feature was the median of its regional levels. Within-region heterogeneity was the median of its regional IQRs, and between-region heterogeneity was the IQR of its regional levels. These summaries gave nine features (S3 Table).

Session count, number of good units, number of regions, number of distinct probe labels, and laboratory explained up to 38% of the variance of single neural features, so we residualized the neural features on these covariates before PCA. The probe-label count was computed from the distinct probe labels of each mouse rather than from its number of probe insertions. The primary neural representation used the six features derived from firing rate and waveform duration. Amplitude can depend on recording geometry and unit isolation, so a second representation that also included the three amplitude features served as a sensitivity analysis. We also examined a combined representation, in which the behavioral and neural component scores were concatenated after scaling each block to contribute equal total variance.

#### Association with model evidence

We analyzed five candidate signals: standard Q-learning RPE, asymmetric RPE, absolute surprise, the outcome-change signal, and the choice-confidence residual. We defined each mouse’s score as the change in approximate PsyTrack log model evidence relative to the stimulus reference, scaled to 1,000 trials. We used Spearman rank correlations to relate these scores to the behavioral, neural, and combined component scores, with Benjamini–Hochberg correction applied separately within each phenotype family. We assessed the stability of the association between evidence for the choice-confidence residual and the fourth behavioral component using 20,000 permutations, 10,000 mouse-level bootstrap samples, and leave-one-mouse-out analysis. These checks were performed after the association was identified and assessed its stability within the same cohort.

### Ground-truth simulations

The preceding analyses measured how much model-derived signals contributed to predicting recorded choices. To assess what those comparisons could reveal about the underlying decision rule, we simulated two populations with known choice mechanisms. One population used recent trial events directly. The other updated action values using a reward prediction error. We used these populations to test whether a lag-one RPE could improve prediction for a history-based agent and whether a multi-trial RPE trace improved prediction for an agent whose choices depended on accumulated prediction errors.

Each population comprised 60 simulated mice, each completing four sessions of 600 trials in a simplified visual choice task. Signed contrasts were sampled uniformly from the list [*±*1, *±*0.25, *±* 0.125, *±*0.0625, 0, 0], with zero represented twice. The correct side was assigned randomly on zero-contrast trials. We used the same contrast transformation as in the behavioral analyses, Eq (2). The simulations omitted the blockwise stimulus prior.

For the history-based agent, the probability of choosing right was *P* (*c*_*t*_ = +1) = *σ*[*b* + *w*_*s*_*f* (*s*_*t*_) + *h*_*c*_*c*_*t−*1_ + *h*_*w*_*c*_*t−*1_(2*r*_*t−*1_ *−* 1) + *gf* (*s*_*t−*1_)] , (16) where *σ* is the logistic function. The history-based agent’s parameters were drawn from *b ∼U* (*−* 0.3, 0.3), *w*_*s*_*∼ U* (3, 6), *h*_*c*_, *h*_*w*_ *∼U* (0, 0.8), and *g ∼U* (*−*0.8, 0.8).

For the RPE-learning agent, the probability of choosing right was

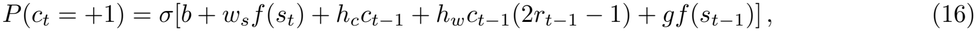

After observing the outcome, the agent updated the chosen action value by *α δ*_*t*_, where

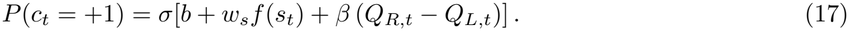

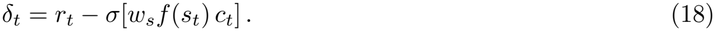

The second term is the agent’s stimulus-based probability that its chosen action was correct. After the update, both action values decayed toward 0.5. The value weight *β*, learning rate *α*, and decay fraction were drawn from *U* (1, 4), *U* (0.2, 0.8), and *U* (0.02, 0.2), respectively. Because *δ*_*t*_ does not depend on the action values, the difference *Q*_*R,t*_ *−Q*_*L,t*_ is an exponentially weighted sum of the past values of *α c δ*, with retention equal to one minus the decay fraction (0.80 to 0.98). This agent’s decision variable is therefore itself a trace of a choice-aligned, stimulus-dependent prediction error.

We analyzed both populations using leave-one-session-out logistic regression with fixed regularization (*C* = 1) and predictor standardization fitted on the training sessions. For the state RPE used in the fitted choice models, we estimated expected reward from smoothed stimulus-by-choice reward rates in the training sessions. We first added the lag-one state RPE to reference models that successively included one-trial choice and reward history, previous stimulus and state terms, and exponential traces of observable history. We then added a state RPE trace to the reference that already contained observable-history traces, using the same fixed retention, *λ* = 0.7 in Eq (12), for both. Unlike the analyses of mouse data, the simulations used fixed regularization, a fixed retention, a tabulated reward expectation, and no block prior. For each simulated mouse, we expressed the change in held-out log likelihood per 1,000 trials and evaluated population effects with two-sided Wilcoxon signed-rank tests.

### Ethics statement

This study analyzed only publicly available data and involved no new experiments on animals. The experiments that produced the data were performed by the International Brain Laboratory under the institutional and national approvals described in the original publications [10, 11].

## Statistical analysis

For each pair of choice models, we summed the held-out log likelihoods across sessions within each mouse. For mouse *m*, we expressed the difference between the augmented model and its matched reference per 1,000 held-out trials,

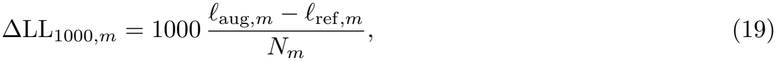

where *l*_aug,*m*_ and *l*_ref,*m*_ are the summed held-out log likelihoods of the augmented and reference models, respectively. *N*_*m*_ is the number of held-out trials for that mouse. Positive values indicate that the augmented model assigned higher probability to the observed choices.

We used the mouse as the unit of inference. We summarized paired effects by their median across mice and obtained 95% percentile confidence intervals using 10,000 bootstrap resamples of mice with replacement. We tested paired differences using two-sided Wilcoxon signed-rank tests. We applied Holm correction separately across the eight one-trial comparisons, the eight recorded-block comparisons, the six multi-trial comparisons, and the three GLM-HMM comparisons. Fixed-retention sensitivity tests formed a separate correction family. For the primary matched-trace comparison, evidence of an incremental RPE contribution required a positive median effect, a bootstrap confidence interval entirely above zero, and a Holm-adjusted *p <* 0.05.

For the phenotype analyses, we used Spearman rank correlations and applied Benjamini–Hochberg correction separately within the behavioral, neural, and combined phenotype families.

## Results

### An exploratory screen of candidate signals

Absolute surprise was the only candidate with a positive median change in approximate PsyTrack log model evidence across 139 mice (S4 Fig). It increased the evidence in 78 of 139 mice, compared with 25 of 139 for the standard Q-learning RPE. This screen compared models fitted to the same trials. The subsequent analyses assessed predictive performance on held-out sessions. Because each candidate was multiplied by the previous choice, the initial evidence increase may partly reflect information about the previous choice itself. In the held-out comparisons, absolute surprise provided little additional predictive information once the reference included the previous choice and reward (Table 4A).

**Table 4.** Held-out choice-model comparisons.

| Added to model | Reference model | Median | 95% CI | Holm *p* |
|---|---|---|---|---|
| *A. One-trial comparisons* |  |  |  |  |
| Previous choice | Current stimulus | 32.008 | [27.473, 35.206] | < 0.001 |
| Choice and reward history | Current stimulus | 30.634 | [26.853, 34.675] | < 0.001 |
| Absolute surprise | One-trial history | 0.008 | [−0.192, 0.140] | 0.716 |
| Signed Q-learning RPE | One-trial history | −0.365 | [−0.682, −0.052] | 0.153 |
| State RPE | One-trial history | 0.596 | [0.412, 0.933] | < 0.001 |
| Previous stimulus terms | One-trial history | 1.328 | [0.912, 1.945] | < 0.001 |
| State RPE | History + previous stimulus | −0.064 | [−0.148, 0.008] | 0.716 |
| State expectation | History + previous stimulus | −0.064 | [−0.122, 0.009] | 0.716 |
| *B. Multi-trial comparisons* |  |  |  |  |
| Discrete lags 2–3 | One-trial state | 16.927 | [14.414, 19.372] | < 0.001 |
| Observable-history traces | One-trial state | 22.713 | [19.663, 25.624] | < 0.001 |
| Lag-one state RPE | State + discrete lags | −0.064 | [−0.161, 0.001] | 0.390 |
| State RPE trace | State + discrete lags | 0.173 | [−0.060, 0.381] | 0.099 |
| Lag-one state RPE | State + history traces | −0.088 | [−0.149, −0.010] | 0.386 |
| Matched state RPE trace | State + history traces | 0.056 | [−0.071, 0.260] | 0.390 |

Values are median changes in held-out log likelihood per 1,000 trials across 107 mice. Positive values favor the model with the added predictor or predictors. Confidence intervals are 95% mouse-level percentile bootstrap intervals. Two-sided Wilcoxon signed-rank *p*-values were adjusted by Holm within each panel. All models include the current stimulus. One-trial history includes the preceding choice, centered reward, and their interaction. The previous-stimulus terms follow the specification of each analysis in Methods. Panel B uses the one-trial state reference without the separate preceding left and right contrast terms (Table 3). History traces summarize choice, centered reward, and their interaction. The matched RPE trace uses the same retention parameter as the observable-history traces. Small negative gains can arise when an additional predictor increases estimation variability without contributing useful predictive information. A negative estimate alone does not establish redundancy. The bootstrap confidence intervals are unadjusted for multiple comparisons, whereas the reported *p*-values are Holm-adjusted.

### Recent trial history improves held-out choice prediction

Fig 2 and Table 4A summarize the one-trial held-out comparisons in 107 mice (427 sessions, 275,292 trials).

**Fig 2.**
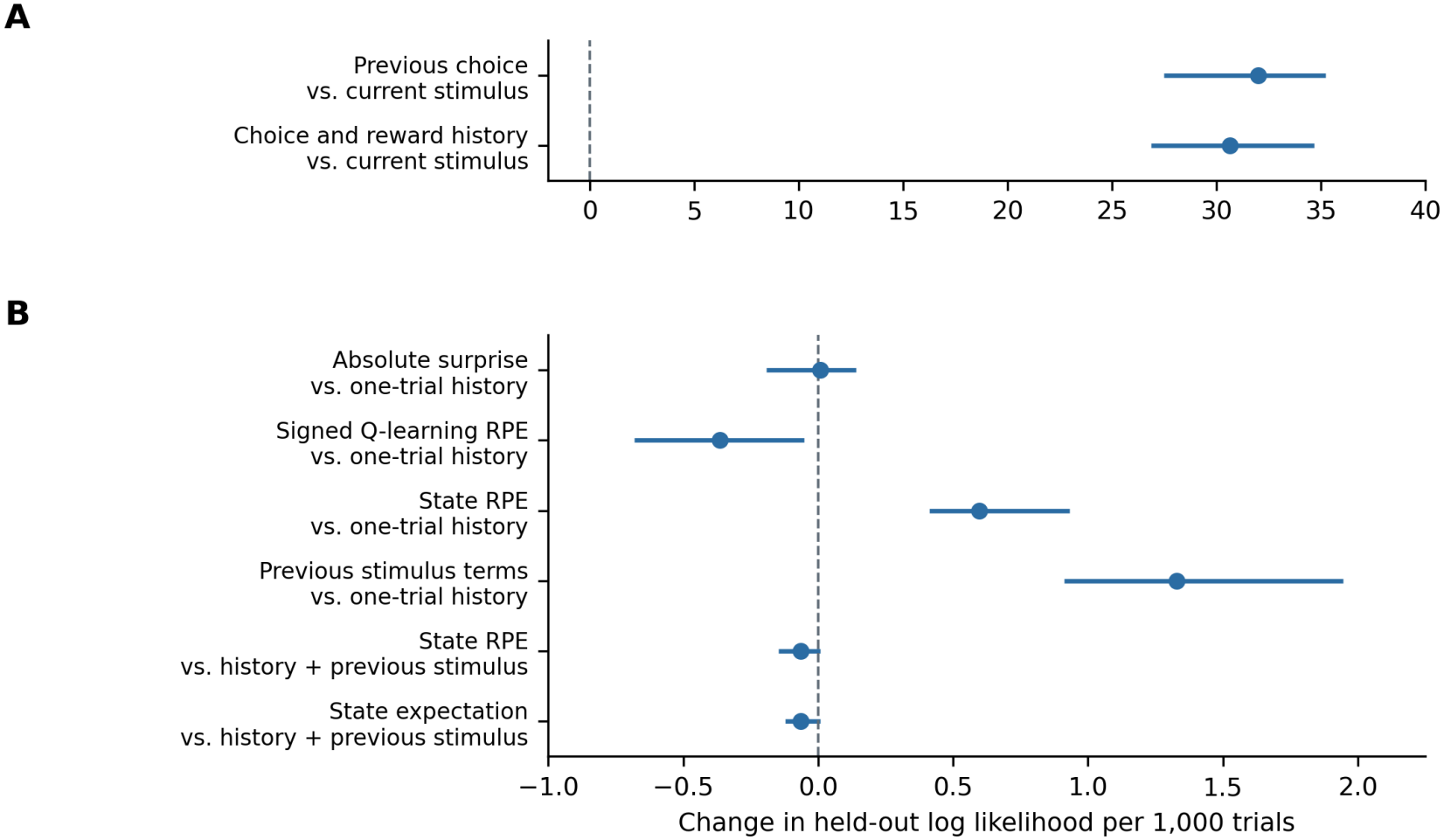
One-trial history and state RPE in held-out choice prediction. Each row adds the named predictor or predictors to the reference model named below it. (A) Additions to the stimulus-only model. (B) Additions to the one-trial history reference, with or without the previous-stimulus terms, on an expanded scale. Points show median changes in held-out log likelihood per 1,000 trials across 107 mice. Horizontal bars show 95% mouse-level bootstrap intervals. The dashed line marks zero gain.

Adding the preceding choice, reward, and their interaction to the stimulus-only model improved prediction in all 107 mice. The median gain was 30.63 log-likelihood units per 1,000 trials (95% bootstrap CI 26.85 to 34.68). The preceding choice alone also improved prediction in all 107 mice, with a median gain of 32.01 units (95% bootstrap CI 27.47 to 35.21). These two medians come from separate comparisons with the stimulus-only model, so their difference does not estimate the effect of adding the preceding reward to the preceding choice. An additional paired comparison outside the main correction family assessed the contribution of reward beyond the preceding choice. Adding the preceding reward and the choice-by-reward interaction produced a median gain of 0.12 log-likelihood units per 1,000 trials (95% bootstrap CI *−*0.32 to 0.94; 56 of 107 mice improved; unadjusted Wilcoxon *p* = 0.25). Much of the predictive improvement over the stimulus-only model was therefore already obtained from the preceding choice. In a separate analysis with fixed *C* = 1, absolute surprise also improved prediction over the stimulus-only model in all 107 mice, with a median gain of 14.85 units. To assess its contribution beyond behavioral history, the next comparisons included the preceding choice, reward, and their interaction in the reference model.

### State RPE adds a small gain beyond choice and reward history

Table 4A and Fig 2 show the effect of adding each RPE-derived predictor to the one-trial history reference. Adding the signed Q-learning RPE gave a median change of *−*0.36 log-likelihood units per 1,000 trials, and adding absolute surprise gave a median change of 0.01 units. Neither predictor showed evidence of improved held-out prediction beyond the one-trial history reference (Table 4A).

The state RPE, whose reward expectation was estimated from the preceding stimulus and choice, improved held-out prediction in 81 of 107 mice. The median gain over the one-trial history reference was 0.60 log-likelihood units per 1,000 trials (95% bootstrap CI 0.41 to 0.93). Because the state RPE incorporates the preceding stimulus, the next comparison tested its contribution after including that stimulus in the reference model.

### The state RPE adds little after including the previous stimulus

The last rows of Table 4A and Fig 2 add the preceding stimulus to the reference. Adding preceding-stimulus terms to the one-trial history model improved prediction by a median of 1.33 log-likelihood units per 1,000 trials. With those terms in the reference, the median change from adding the state RPE was *−*0.064 (95% bootstrap CI *−*0.148 to 0.008). The median change from adding the choice-aligned reward expectation was also near zero. A separate held-out analysis gave a similar pattern across mice (S5 Fig). The state RPE therefore provided little additional predictive information once the reference included both one-trial history and the preceding stimulus.

Controlling for the recorded current and preceding block probabilities produced a similar pattern (S1 Fig). With these controls, the state RPE added a median of 0.45 log-likelihood units per 1,000 trials over one-trial history. Including the preceding stimulus reduced the median change to *−*0.13 (95% bootstrap CI *−*0.18 to *−*0.01). The next comparison asked whether an RPE trace added predictive value when the reference also summarized earlier choices and rewards.

### Multi-trial history improves prediction with little additional gain from the RPE trace

Adding choice and reward history from two and three trials back improved prediction over the one-trial state model (Table 4B). Observable-history traces also improved prediction over that model, with a median gain of 22.71 log-likelihood units per 1,000 trials (95% bootstrap CI 19.66 to 25.62, Fig 3A).

**Fig 3.**
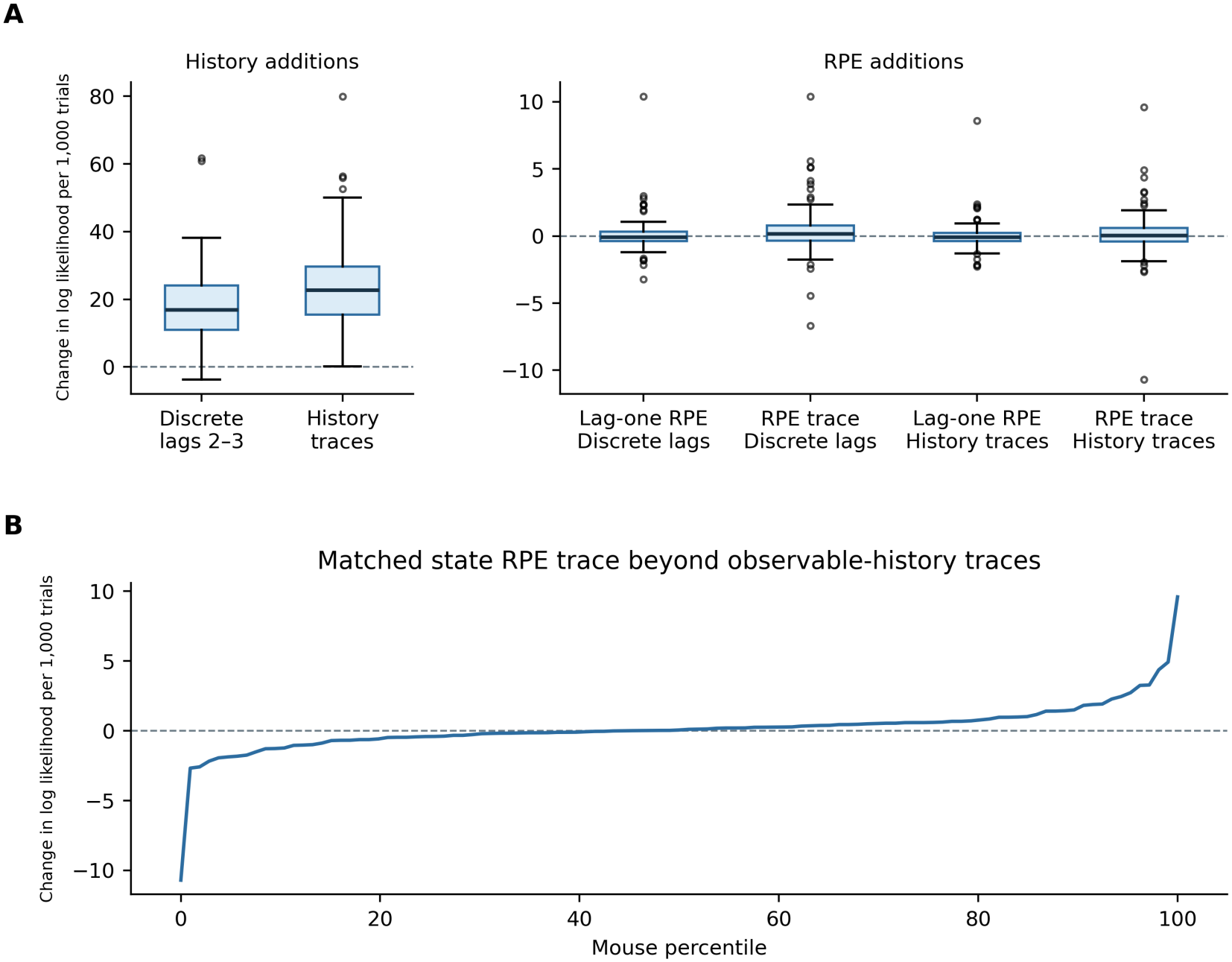
Multi-trial history and the matched state RPE trace. (A) Distributions of mouse-level held-out gains for all six multi-trial comparisons. History additions and RPE additions use separate vertical scales. The labels below each RPE addition identify its multi-trial reference. The first two comparisons add discrete lags or observable-history traces to the one-trial state model. The remaining comparisons add the lag-one state RPE or its trace to each multi-trial reference. (B) Mouse-level gains for the primary comparison, ordered by percentile. This comparison adds the state RPE trace to a model with observable-history traces at the same retention parameter. Both panels show changes in held-out log likelihood per 1,000 trials across 107 mice. In panel A, boxes span the interquartile range and lines mark medians. Whiskers extend to the most extreme values within 1.5 interquartile ranges of each box, and points mark values beyond the whiskers.

Adding the state RPE trace to the observable-history trace model gave a median change of 0.056 log-likelihood units per 1,000 trials (95% bootstrap CI *−*0.071 to 0.260, Fig 3B). For scale, the upper confidence limit was approximately 1% of the median gain obtained by adding observable-history traces.

To assess whether the result depended on the tested retention range, the comparison was repeated with a wider grid. With this grid, 215 of 427 folds (50%) selected a retention above 0.70, the largest value of the original grid, and 25 folds selected the new largest value, 0.98. With this grid, the median change from the matched state RPE trace was 0.036 (95% bootstrap CI *−*0.109 to 0.267; 56 of 107 mice improved; S2 Fig). The fixed-retention tests yielded little evidence for a gain, and prediction worsened at the highest retention tested (median *−*0.42 at *λ* = 0.98, S2 Fig). Once the reference model included observable-history traces with the same retention, adding the state RPE trace did not reliably improve held-out prediction.

### A GLM-HMM improves prediction beyond the fixed three-lag model

Fig 4 compares GLM-HMMs with the fixed three-lag model on the same held-out folds.

**Fig 4.**
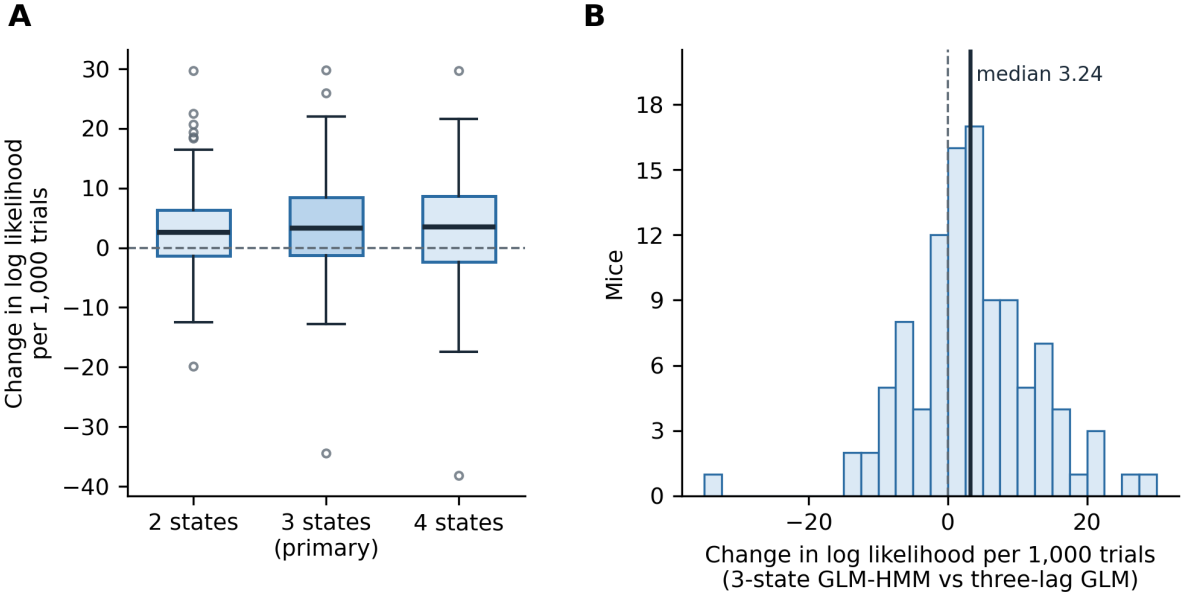
Held-out prediction with a GLM-HMM. (A) Distributions of held-out log-likelihood gains per 1,000 trials across 107 mice for two-, three-, and four-state GLM-HMMs compared with the fixed three-lag GLM. Boxes span the interquartile range and lines mark medians. Whiskers extend to the most extreme values within 1.5 interquartile ranges of each box, and points mark values beyond the whiskers. The dashed line marks zero gain. (B) Histogram of mouse-level gains for the primary three-state comparison. The dashed vertical line marks zero gain and the solid vertical line marks the median.

The three-state GLM-HMM improved held-out log likelihood over the fixed three-lag model in 73 of 107 mice. The median gain was 3.24 log-likelihood units per 1,000 trials (95% bootstrap CI 2.19 to 4.05, Fig 4). Two- and four-state models also showed positive median gains. The gain in log likelihood was accompanied by little change in median classification accuracy. Many fits reached the iteration limit before meeting the convergence criterion, which limits interpretation of the estimated gain. The next analysis examined whether between-mouse differences in candidate-signal model evidence were associated with behavioral and neural phenotypes.

### Model evidence for the choice-confidence residual covaries with behavioral PC4

After adjustment for session count, laboratory, and task protocol, 11 behavioral principal components explained 81% of the variance in the adjusted behavioral features. The fourth behavioral component (PC4) was positively associated with the gain in PsyTrack model evidence from adding the choice-confidence residual (Spearman *ρ* = 0.286, adjusted *q* = 0.036, Fig 5). This was the only behavioral association to survive correction across 55 signal–component comparisons, and it remained positive when each mouse was omitted in turn. Absolute surprise showed an association with PC4 in the same direction that did not survive correction (*ρ* = 0.237, *q* = 0.138). No association with a neural component survived correction (largest |*ρ*| = 0.19, *q* = 0.54), and neither did any association with a combined component (largest |*ρ*| = 0.26, *q* = 0.19).

**Fig 5.**
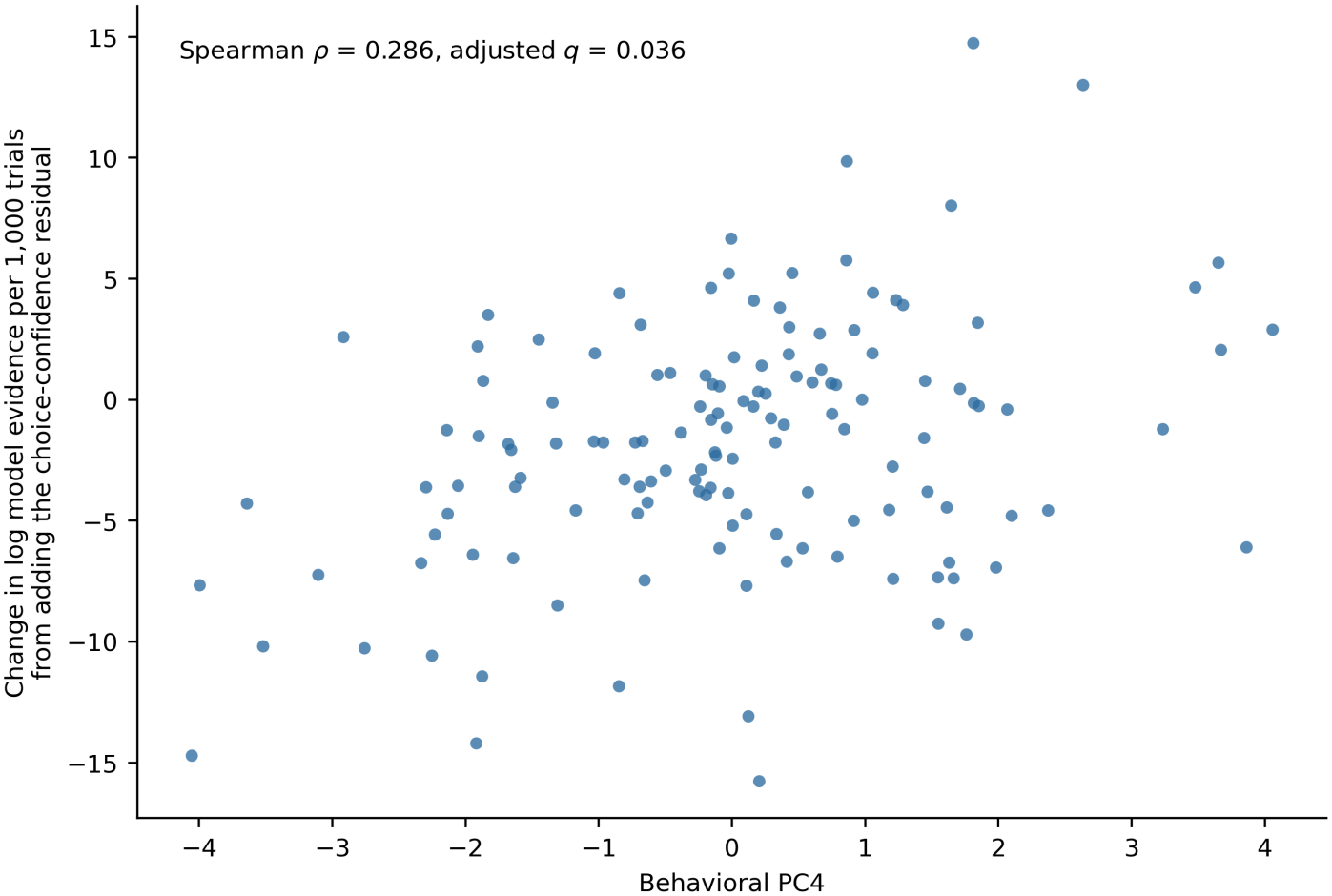
Behavioral PC4 and evidence for the choice-confidence residual. Each point represents one of 139 mice. The horizontal axis shows the fourth behavioral principal component score. The vertical axis shows the change in approximate PsyTrack log model evidence per 1,000 trials when the choice-confidence residual was added to the stimulus reference. The plot reports the Spearman correlation and its false-discovery-rate-adjusted *q*-value.

Mice with higher PC4 scores tended to show larger pupil responses to the go cue and reward, greater sensitivity to the block prior, more choice repetition and omissions, and less wheel activity. Selected PC4 loadings and robustness estimates are provided in S1 Table.

### The matched state RPE trace improves prediction for the simulated learner

The simulated agents tested how the held-out comparisons behaved when choices followed known history-based or RPE-based decision rules (Fig 6).

**Fig 6.**
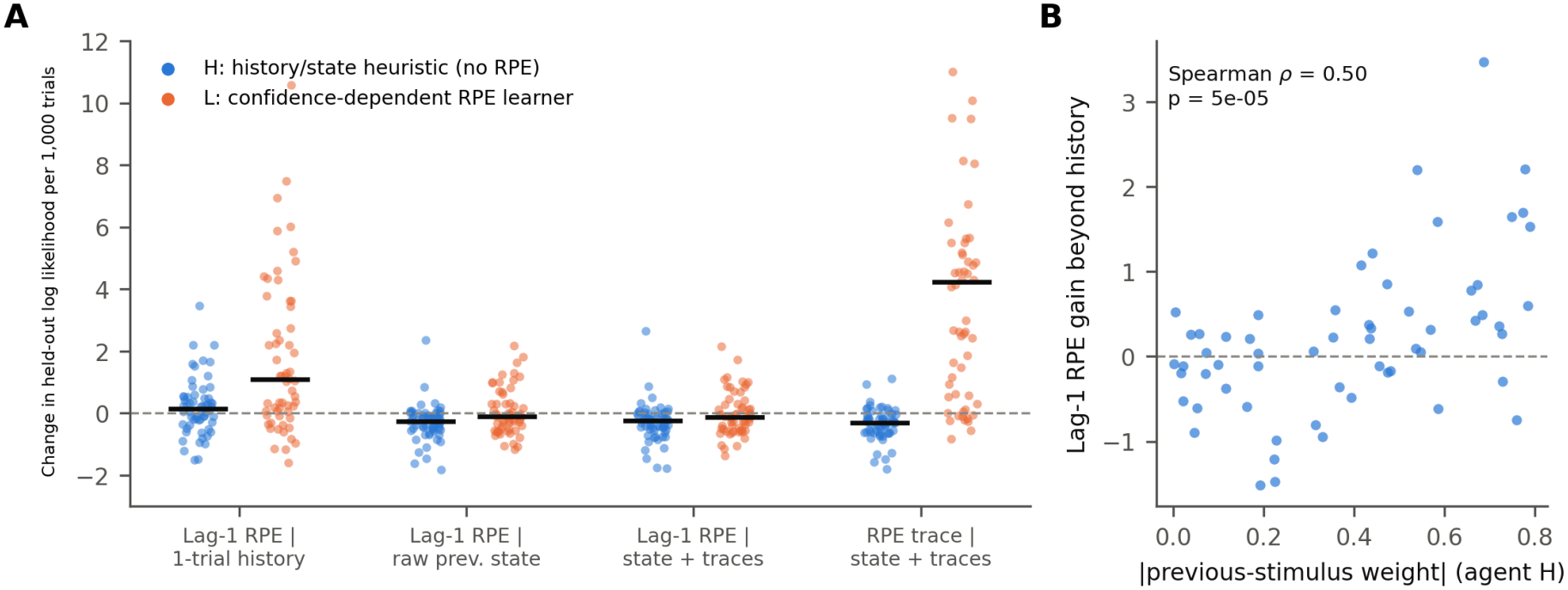
State RPE effects for agents with known choice rules. (A) Changes in held-out log likelihood per 1,000 trials for 60 simulated mice per agent type. The first three comparisons add the lag-one state RPE to one-trial history, previous state, and previous state with observable-history traces. The fourth adds a state RPE trace to the state reference with observable-history traces at the same retention parameter. Points show simulated mice and black horizontal marks show medians. (B) In the history-based agent, the lag-one state RPE gain relative to the one-trial history model is shown against the magnitude of the preceding-stimulus weight used to generate choices.

In the simulated history-based agent, adding the lag-one state RPE to the one-trial history model gave a median gain of 0.15 log-likelihood units per 1,000 trials (*p* = 0.16). The gain was larger in agents whose choices depended more on the preceding stimulus (Spearman *ρ* = 0.50, Fig 6B). With the preceding stimulus included directly, the median change was *−*0.25 (Fig 6A).

For the simulated RPE learner, adding the lag-one state RPE gave little gain after the reference included the preceding stimulus. The matched state RPE trace improved prediction by a median of 4.22 log-likelihood units per 1,000 trials over the reference model containing observable-history traces with the same retention parameter. The corresponding median change for the history-based agent was *−*0.30. The simulated learner’s decision variable had the same form as the tested predictor (Methods). This result therefore demonstrates sensitivity to the specified RPE-based choice rule.

## Discussion

The incremental predictive value of the state RPE depended on how the reference model represented earlier trials. Including the preceding stimulus brought the lag-one gain close to zero. Extending observable choice and reward history over several trials improved prediction, and the matched state RPE trace added little over those history traces.

The choice-aligned reward term of the state RPE is already represented by the observable-history predictors (Eq (10) and Eq (13)). The remaining term is a fitted reward expectation that depends on stimulus and choice, and this term can supply information about the preceding stimulus. The lag-one state RPE can therefore improve prediction when the reference omits the preceding stimulus. At the trace level, the comparison assesses the additional predictive contribution of the choice-aligned reward expectation beyond choice and reward traces with the same retention. In mouse perceptual tasks, past confidence and reward influence later choices [16], and those choices can reflect events over several trials [7].

The simulations illustrate how these comparisons behave under two specified choice rules. In the history-based agent, the apparent gain from the lag-one state RPE increased with the weight assigned to the preceding stimulus. This pattern shows how an RPE predictor can capture omitted stimulus information when choices are generated directly from trial history. In the RPE learner, the lag-one state RPE gave little gain after the reference included the preceding stimulus, whereas the matched state RPE trace improved prediction. The simulated learner accumulated a choice-aligned state RPE, matching the form of the tested trace. Its positive result therefore establishes sensitivity under this specified choice rule. Sensitivity to other learning rules, including Q-learning with value-dependent errors, and to smaller effects remains untested. The size of the gain also depends on the parameter ranges simulated here.

The GLM-HMM improved prediction beyond the fixed three-lag model, indicating that the fixed coefficients and explicit history terms did not capture all predictive structure. This is consistent with evidence that mice shift among decision strategies during perceptual tasks [18]. In our analysis, the GLM-HMM and the fixed model received the same explicit predictors. Because the GLM-HMM updates its state probabilities throughout a session, the inferred state could also carry information from choices earlier than the three explicit lags. The gain may therefore reflect changes in how mice use the stimulus and recent history, dependence on earlier choices, or both. Most three-state fits had not met the convergence criterion when optimization ended, so the estimated gain may change with further optimization.

Across mice, the gain in PsyTrack model evidence from the choice-confidence residual covaried with behavioral PC4. The component combined variation in pupil responses, prior sensitivity, choice repetition, omissions, and wheel activity. Pupil responses track arousal-related changes in cortical state [22, 23], but whether arousal-related changes underlie the association observed here remains untested. The choice-confidence residual and several PC4 features, including prior sensitivity and choice repetition, were computed from the same choices. Their shared dependence on choice behavior may therefore contribute to the association. The association concerns the composite behavioral component and does not identify which features account for it. Its generalizability remains to be tested in independent data.

Dopamine recordings and manipulations have shown that some midbrain responses reflect the difference between received and expected reward [1, 2]. In a different cue–reward paradigm, mesolimbic dopamine release tracked retrospective associations between rewards and the cues that preceded them [26]. Our analysis assessed whether model-derived RPE signals contributed to choice prediction beyond observable history and stimulus information. The limited incremental contribution therefore constrains their predictive value in this task, without determining what dopamine signals encode.

Our conclusions apply to one-step-ahead choice prediction in the IBL visual decision task. We estimated reward expectation from stimulus and choice, and represented longer history using discrete lags and exponential traces. The recorded block probability controlled for the stimulus prior programmed by the task. An individual mouse’s belief about the current block could differ from that prior, particularly around block transitions, and could change the reward expectation used to compute an RPE. Dopamine prediction errors can reflect such inferences about hidden task states [27].

The sessions came from trained mice recorded for the Brain-Wide Map. The contribution of value learning may differ during task acquisition, which lies outside the present analysis.

For the matched state RPE trace, the 95% confidence interval for the median gain ranged from *−*0.071 to 0.260 log-likelihood units per 1,000 trials. This estimate constrains the incremental predictive contribution without establishing that it is zero. The parameter-recovery analysis adds a related caution: our simulations suggest that weak sensitivity to action values can make fitted learning rates difficult to interpret (S1 Appendix, S3 Fig), while previous work has shown that omitting choice perseveration can produce spurious learning-rate asymmetries [14].

Recent choices and outcomes may already reflect the behavioral consequences of prediction-error learning. The small gain from the matched state RPE trace therefore limits what these choice-prediction comparisons can reveal about how mice learned from those outcomes.

## Acknowledgments

Claude (Anthropic) and ChatGPT (OpenAI) were used for coding assistance and manuscript revision, including improvements to language, clarity, and organization. The authors reviewed the AI-assisted code and text, checked the analysis outputs, and take responsibility for the final manuscript.

## Data availability

The source trial data are available from the DANDI Archive as IBL Brain Wide Map, Dandiset 000409, version 0.260309.1324 [12]. The phenotype analyses used the IBL data files of the same sessions from the IBL public database through ONE at https://openalyx.internationalbrainlab.org (Table 1). Session identifiers, relative file paths, collections, and revisions are documented in the source-file inventory supplied with S1 Dataset. The supporting dataset is included as ancillary material with this preprint. Analysis code and additional materials are available from the corresponding author, Alireza Khanteymoori, upon request. The accompanying S1 Dataset contains saved mouse-level choice-model results, behavioral and neural feature tables, PC4 association data, and reported simulation summaries. Its file index documents the source of each included table. The derived trial-level dataset (the variables in Table 1A for all 294,938 trials) is also supplied with the supporting data and can be regenerated from the source data with the extraction code.

## Supporting information

## S1 Appendix. Learning-rate recovery

In the 139 mouse fits, 33.1% of standard Q-learning rate estimates and 47.5% of negative-error learning-rate estimates from the asymmetric model were within 0.01 of an optimization bound (S6 Fig). We used parameter-recovery simulations to test when such boundary estimates can arise. We also tested whether a model with separate learning rates for positive and negative prediction errors could estimate different rates when the simulated agent used a single rate or repeated choices without updating action values.

Each fitting condition used 200 simulated mice, with four sessions of 600 trials per mouse. Standard Q-learning agents chose according to the stimulus, a choice bias, and the difference between the two action values, and updated the chosen action value using a single learning rate *α*. We drew *α* from *U* (0.05, 0.95) and the sensitivity to action values *β* from *U*(0, 4) to assess recovery across a range of contributions of learned values to choice. The stimulus weight and choice bias were drawn from *U*(3, 6) and *U*(*−*0.5, 0.5), respectively. Choice-repetition agents used a running choice history *k*_*t*_ in place of the action-value difference. Their probability of choosing right was *σ*[*b* + *w*_*s*_*f* (*s*_*t*_) + *κk*_*t*_], with *k*_*t*+1_ = *k*_*t*_ + *τ*_*k*_(*c*_*t*_ *−k*_*t*_). We drew the weight on choice history *κ* from *U* (0.5, 2) and its update rate *τ*_*k*_ from *U* (0.1, 0.6).

We fitted the standard Q-learning model to choices from the standard learners and the asymmetric model to choices from both agent types. We used maximum likelihood with L-BFGS-B and six random starts per fit, constraining learning rates to [0, 1]. We quantified recovery of the single learning rate using Spearman correlation and median absolute error between the generating and fitted values. For asymmetric fits, we summarized the difference between the fitted rates, 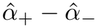 . We also recorded the proportion of estimates within 0.01 of either optimization bound.

### Learning-rate recovery depends on how strongly values guide choice

Among simulated Q-learners, fitted learning rates tracked the generating rates more closely when action values strongly influenced choice. The Spearman correlation between fitted and generating rates was 0.98 for agents with strong value sensitivity and 0.55 for agents with weak value sensitivity. When action values contribute little to choice, changing the learning rate has less effect on the predicted choice probabilities.

Fitting separate learning rates for positive and negative prediction errors also produced apparent asymmetry. The fitted rates differed by more than 0.20 in 35% of simulated learners that used a single rate and in 57% of agents that repeated choices without updating action values. In the choice-repetition group, 49.5% of fitted negative-error rates and 28.5% of fitted positive-error rates were within 0.01 of an optimization bound. The negative-error boundary rate was similar to the 47.5% observed in the mouse fits. These simulations show how weak value sensitivity and omitted choice repetition can make fitted learning rates difficult to interpret.

See S3 Fig. for the plotted recovery and asymmetry summaries.

**S1 Table.** Behavioral PC4 loadings and association checks.

| Measure | Value | Description |
|---|---|---|
| Variance explained | 8.51% | Behavioral PC4 |
| Go-cue pupil response | 0.334 | Selected behavioral PC4 loading |
| Reward pupil response | 0.323 | Selected behavioral PC4 loading |
| No-go fraction | 0.320 | Selected behavioral PC4 loading |
| Prior sensitivity | 0.309 | Selected behavioral PC4 loading |
| Choice repetition | 0.306 | Selected behavioral PC4 loading |
| Wheel active fraction | -0.270 | Selected behavioral PC4 loading |
| Wheel moves per minute | -0.249 | Selected behavioral PC4 loading |
| Number of mice | 139 | Same cohort used for the component and model evidence |
| Spearman rho | 0.285585 | Association with choice-confidence residual evidence |
| Unadjusted p | 0.0006547774119 | Spearman test |
| BH-adjusted q | 0.0360 | Correction within the behavioral association family |
| Permutation p | 0.0009999500025 | 20000 permutations |
| Bootstrap rho lower 95% bound | 0.109761 | 10000 mouse-level bootstrap samples |
| Bootstrap rho upper 95% bound | 0.450925 | 10000 mouse-level bootstrap samples |
| Leave-one-mouse-out rho minimum | 0.270297 | One estimate for each omitted mouse |
| Leave-one-mouse-out rho maximum | 0.303672 | One estimate for each omitted mouse |

**S1 Fig.**
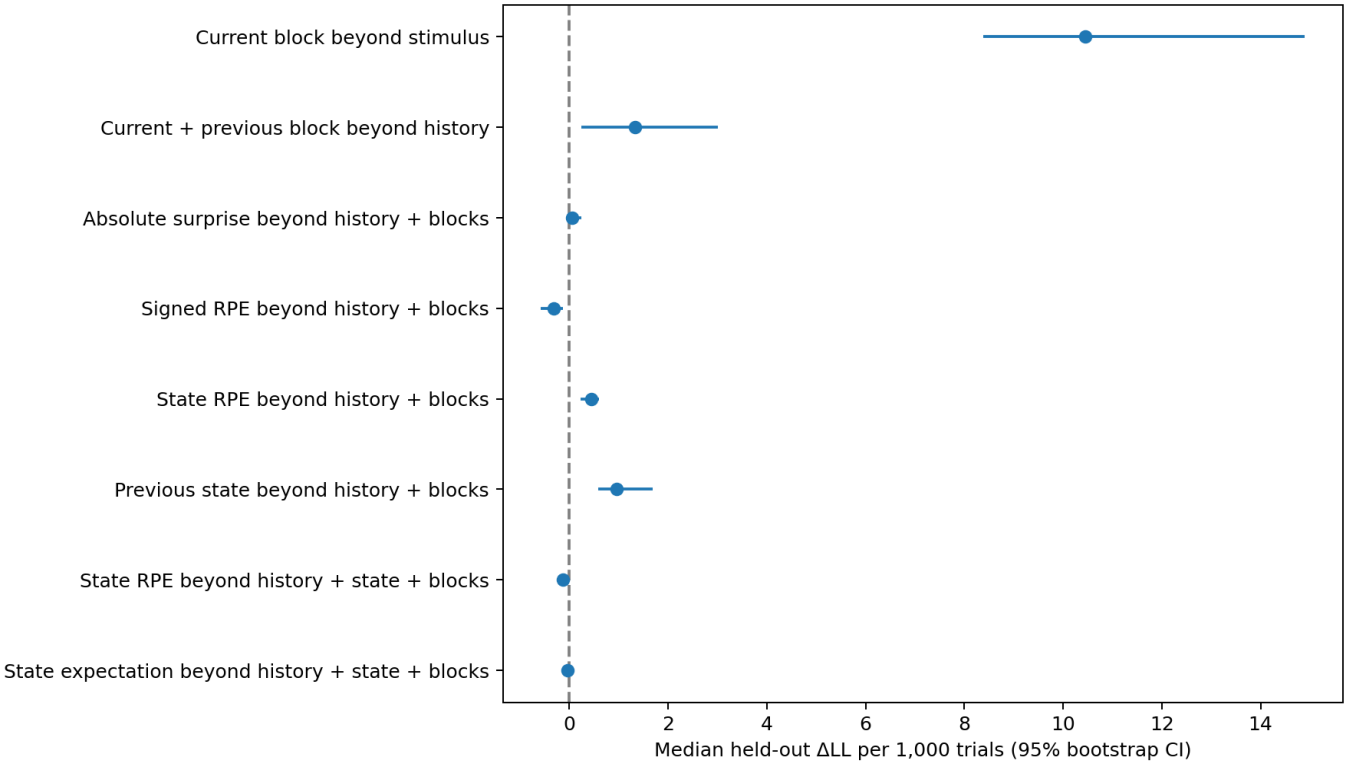
One-trial choice-model comparisons with recorded block controls. The plotted comparisons test the contribution of the recorded current block probability or current and preceding block probabilities, and the incremental effects of preceding stimulus and RPE-derived predictors given the stated reference. Points show medians across 107 mice and bars are 95% mouse-level bootstrap intervals. The dashed line marks zero gain.

**S2 Fig.**
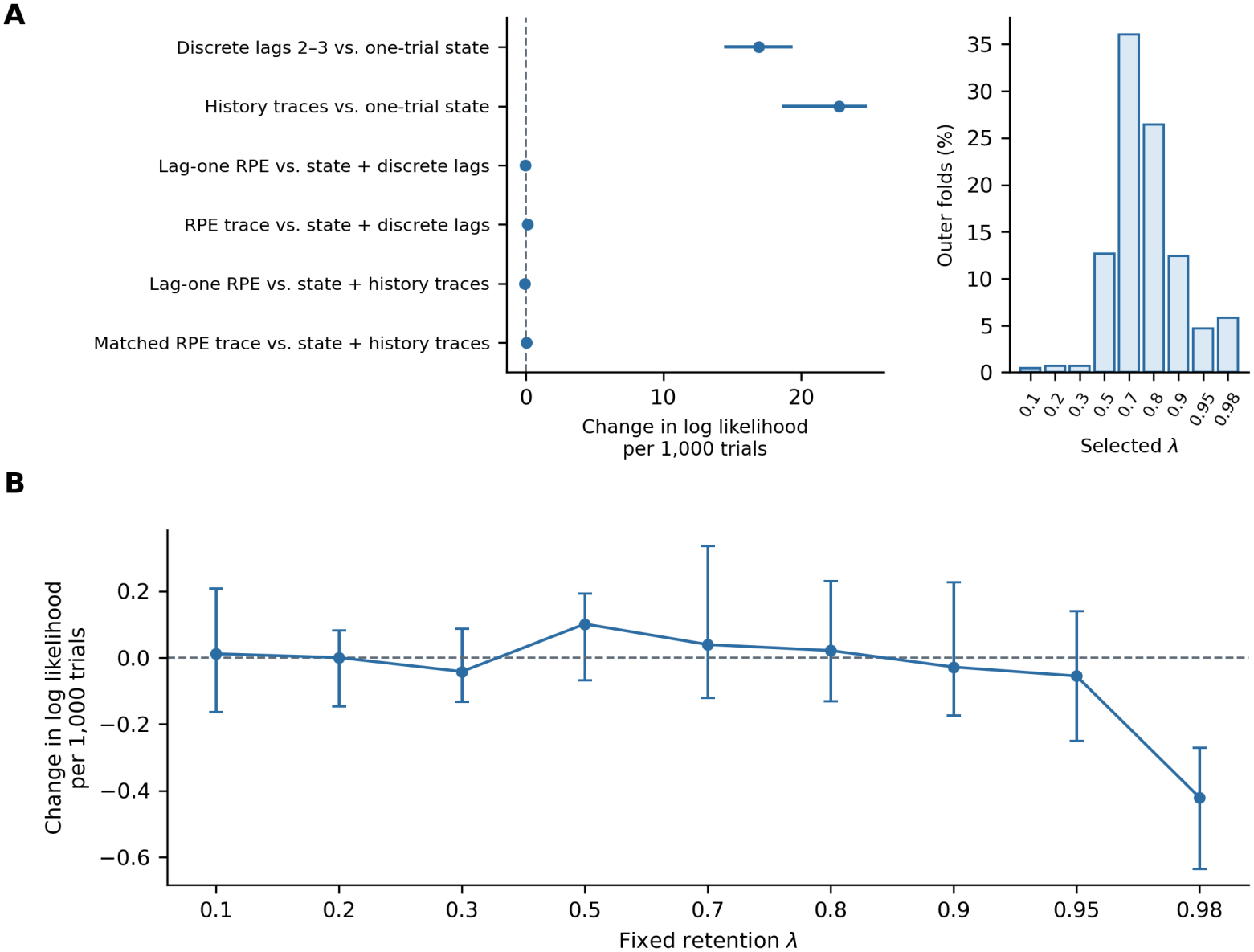
Sensitivity of the matched state RPE trace to retention. (A) Left, median effects and 95% mouse-level bootstrap intervals for the six multi-trial comparisons with the wider selection grid. Right, distribution of training-selected retention values across the 427 outer folds. (B) Change in held-out log likelihood per 1,000 trials from adding the state RPE trace to the observable-history trace model at each fixed retention value of the wider grid. Both traces used the same retention, and the regularization strength was the value selected in the wider-grid analysis. Points show medians across 107 mice and bars show 95% mouse-level bootstrap intervals. The dashed line marks zero change.

**S3 Fig.**
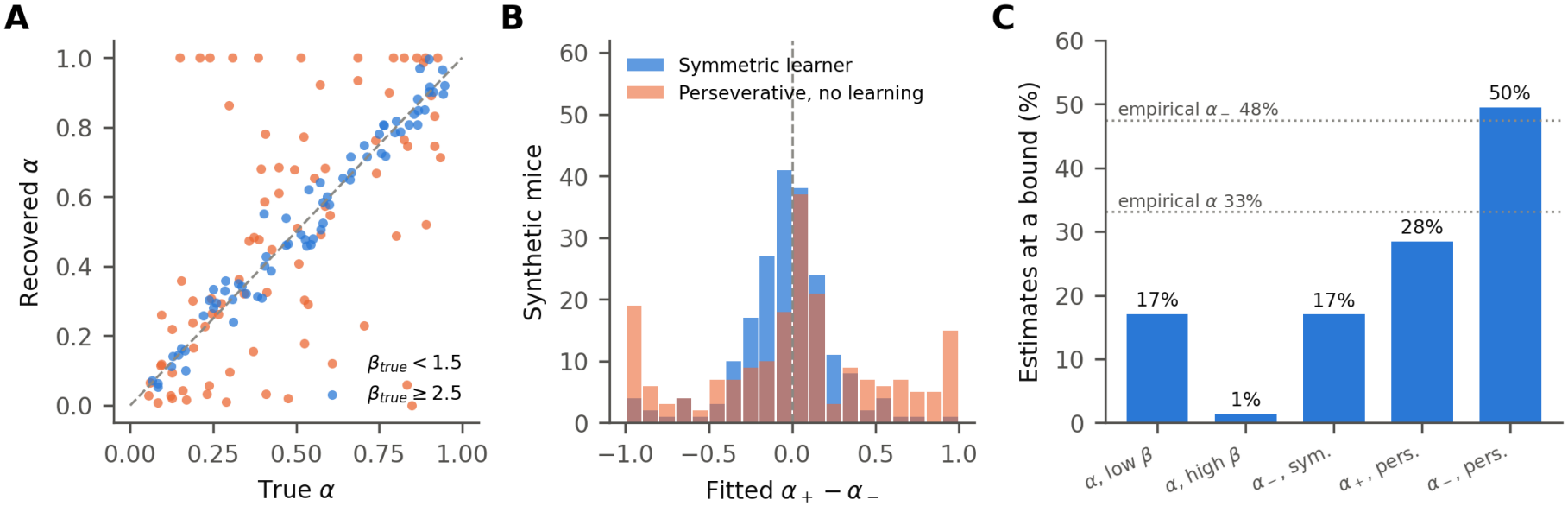
Learning-rate recovery and apparent asymmetry. (A) True and fitted learning rates among standard Q-learners with weak (*β <* 1.5) or strong (*β≥* 2.5) value sensitivity. (B) Fitted differences between positive and negative learning rates for agents generated with a single learning rate or with choice repetition without value updating. (C) Percentages of estimates within 0.01 of an optimization bound. Horizontal dotted lines mark the empirical boundary rates.

**S4 Fig.**
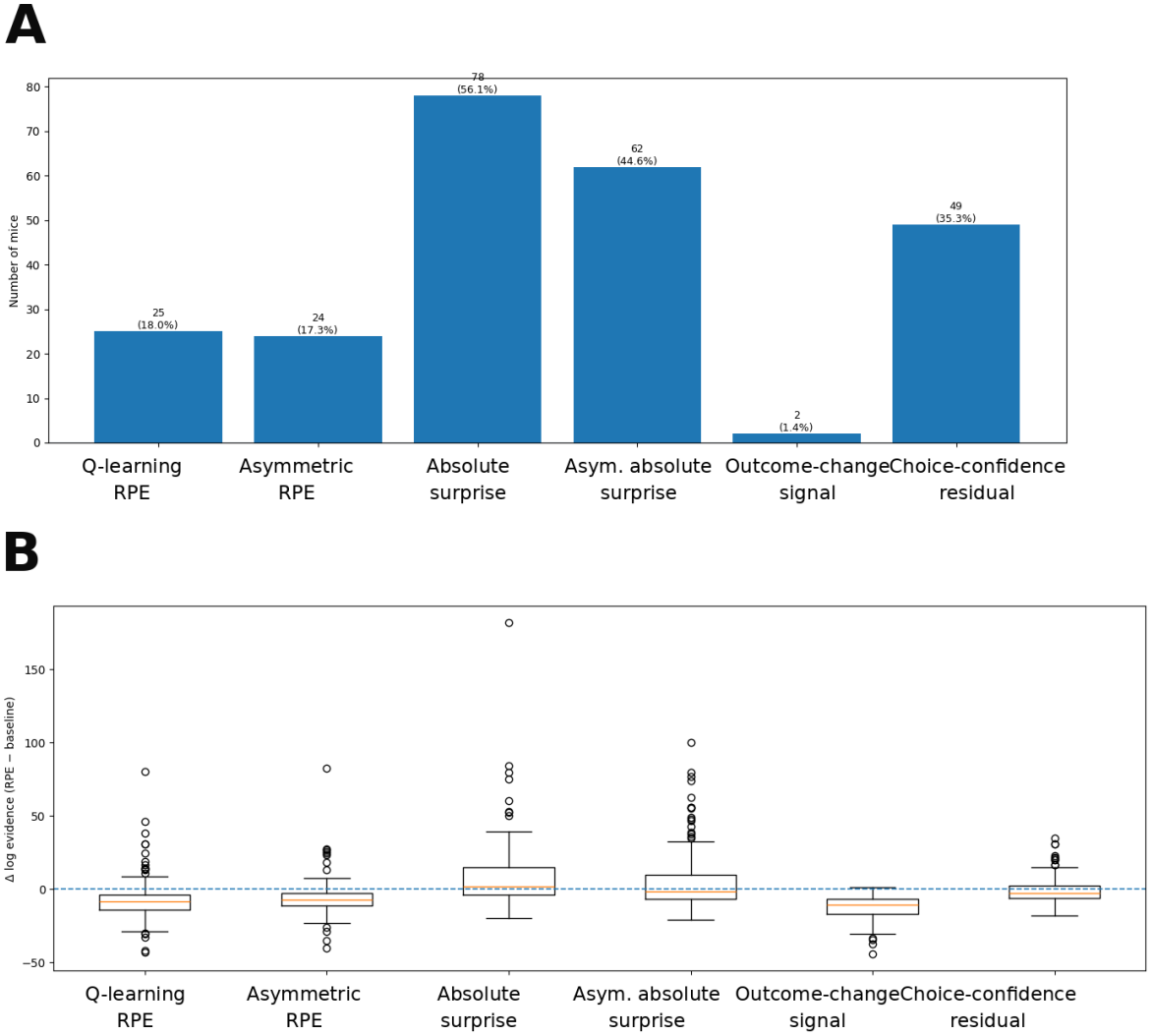
Exploratory PsyTrack screen of six candidate signals. Each candidate was added separately to the same stimulus-only model. (A) Number and percentage of mice with a positive change in approximate log model evidence for each signal. (B) Distribution of evidence changes across 139 mice. The horizontal dashed line marks zero change.

**S5 Fig.**
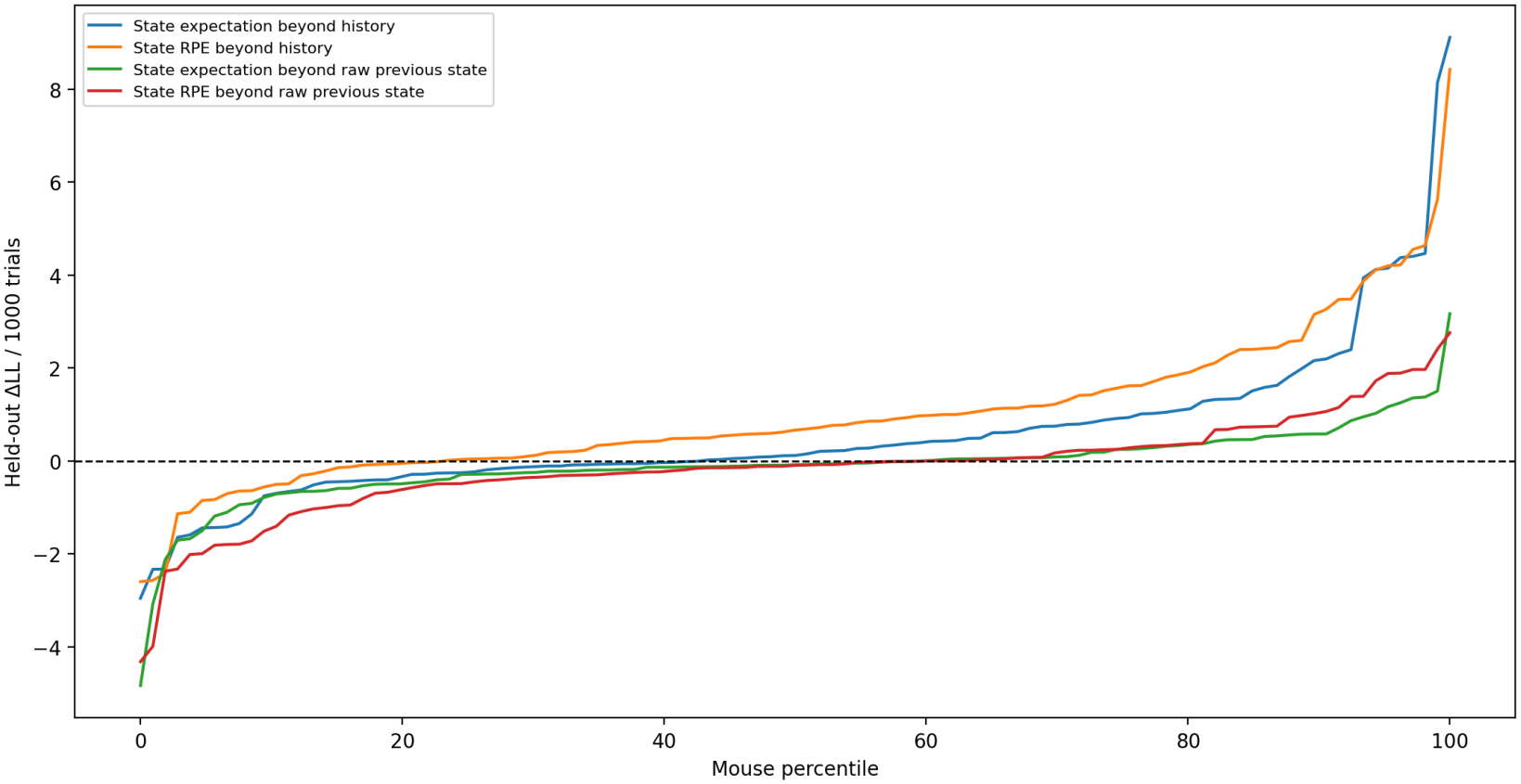
Mouse-level state RPE and state expectation effects with and without previous-state control. In this separate leave-one-session-out analysis, we added either state RPE or state expectation to each of two reference models: choice and reward history alone, or history plus the full previous state. The full previous-state reference included the preceding signed and absolute stimulus, the choice-by-signed-stimulus interaction, and transformed left and right contrasts. We selected L2 regularization strength by inner validation within the training data. Each curve ranks the 107 mice separately by the change in held-out log likelihood per 1,000 trials for that comparison. The dashed line marks zero change.

**S6 Fig.**
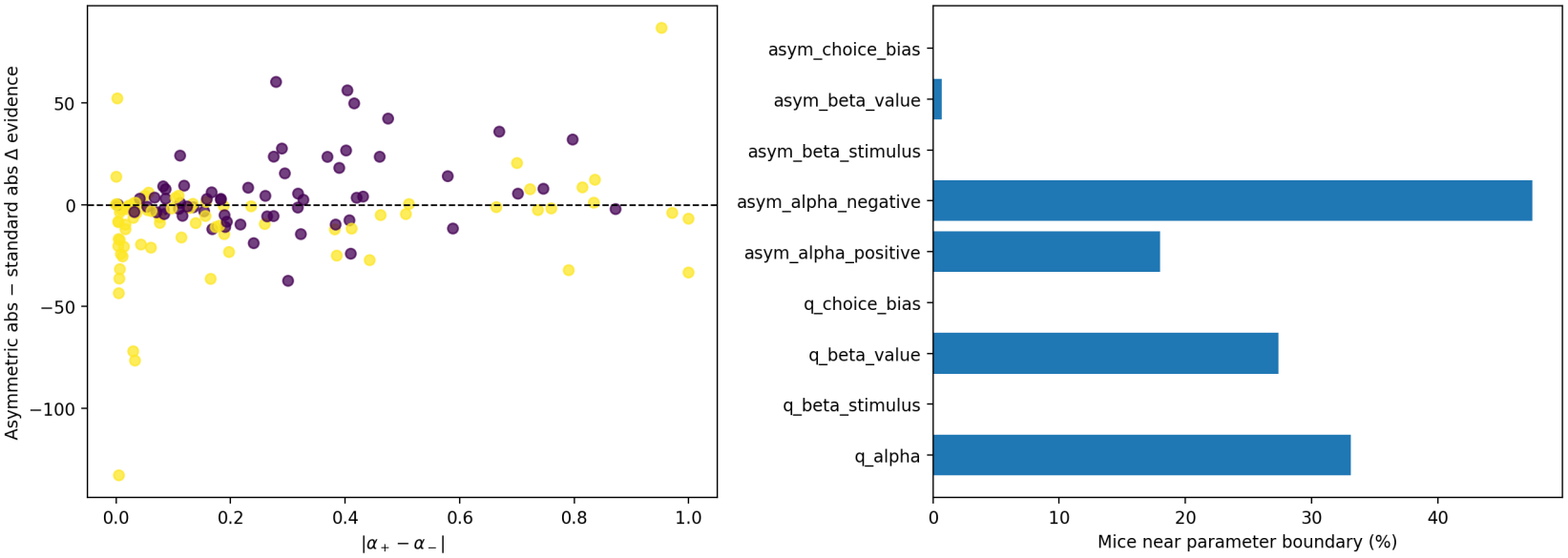
Empirical Q-learning parameter diagnostics. Left, the absolute difference between fitted positive- and negative-error learning rates against the change in exploratory PsyTrack model evidence for asymmetric versus standard absolute surprise (Spearman *ρ* = 0.25, *p* = 0.0029). Each point represents one of 139 mice. Yellow indicates that at least one parameter was flagged as near an optimization bound. Right, percentage of mouse fits flagged near a bound for each parameter.

**S7 Fig.**
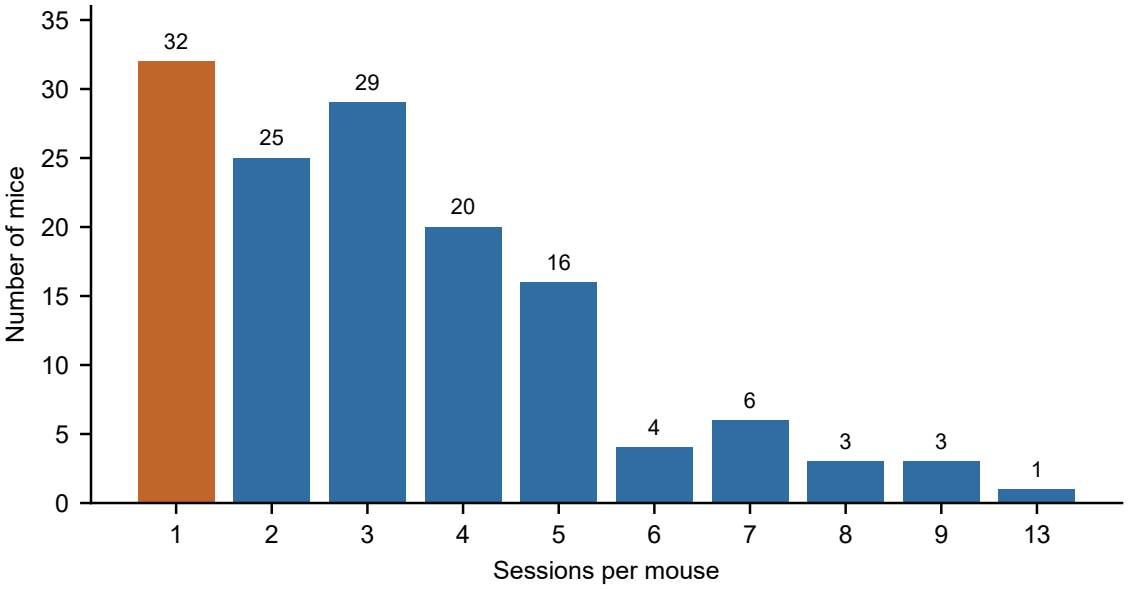
Number of sessions per mouse. Distribution of recorded sessions across the 139 mice (459 sessions). The 32 mice with a single session (orange) were excluded from the held-out analyses, which require at least one training session.

**S2 Table.**
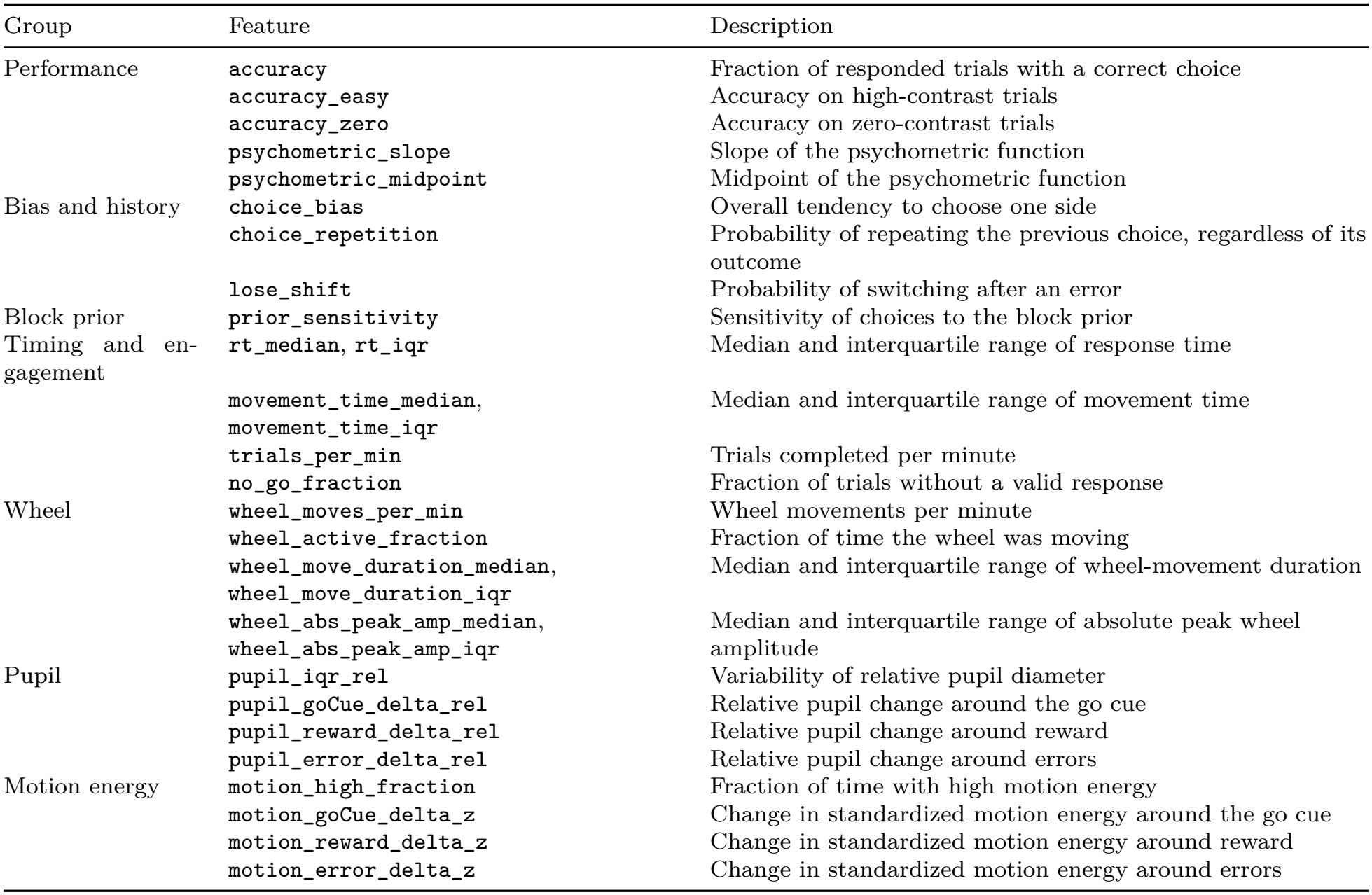
Behavioral features. The 29 mouse-level behavioral features used in the behavioral phenotype. Six further features (win-stay, choice entropy, accuracy on hard trials, median response latency, and pupil and motion responses around feedback) were removed before PCA because each nearly duplicated another feature.

| Group | Feature | Description |
|---|---|---|
| Performance | accuracy | Fraction of responded trials with a correct choice |
|  | accuracy_easy | Accuracy on high-contrast trials |
|  | accuracy_zero | Accuracy on zero-contrast trials |
|  | psychometric_slope | Slope of the psychometric function |
|  | psychometric_midpoint | Midpoint of the psychometric function |
| Bias and history | choice_bias | Overall tendency to choose one side |
|  | choice_repetition | Probability of repeating the previous choice, regardless of its outcome |
|  | lose_shift | Probability of switching after an error |
| Block prior | prior_sensitivity | Sensitivity of choices to the block prior |
| Timing and en-gagement | rt_median, rt_iqr | Median and interquartile range of response time |
|  | movement_time_median, movement_time_iqr | Median and interquartile range of movement time |
|  | trials_per_min | Trials completed per minute |
|  | no_go_fraction | Fraction of trials without a valid response |
| Wheel | wheel_moves_per_min | Wheel movements per minute |
|  | wheel_active_fraction | Fraction of time the wheel was moving |
|  | wheel_move_duration_median, wheel_move_duration_iqr | Median and interquartile range of wheel-movement duration |
|  | wheel_abs_peak_amp_median, wheel_abs_peak_amp_iqr | Median and interquartile range of absolute peak wheel amplitude |
| Pupil | pupil_iqr_rel | Variability of relative pupil diameter |
|  | pupil_goCue_delta_rel | Relative pupil change around the go cue |
|  | pupil_reward_delta_rel | Relative pupil change around reward |
|  | pupil_error_delta_rel | Relative pupil change around errors |
| Motion energy | motion_high_fraction | Fraction of time with high motion energy |
|  | motion_goCue_delta_z | Change in standardized motion energy around the go cue |
|  | motion_reward_delta_z | Change in standardized motion energy around reward |
|  | motion_error_delta_z | Change in standardized motion energy around errors |

**S1 Dataset. Values underlying the figures and tables**. Per-mouse held-out log likelihoods and comparison effects for the one-trial, multi-trial, block-control and GLM-HMM analyses, fixed-retention effects, PsyTrack model-evidence changes, and the available mouse-level behavioral and neural features and component scores. Reported simulation summaries and a source-file index are also included.

**S3 Table.**
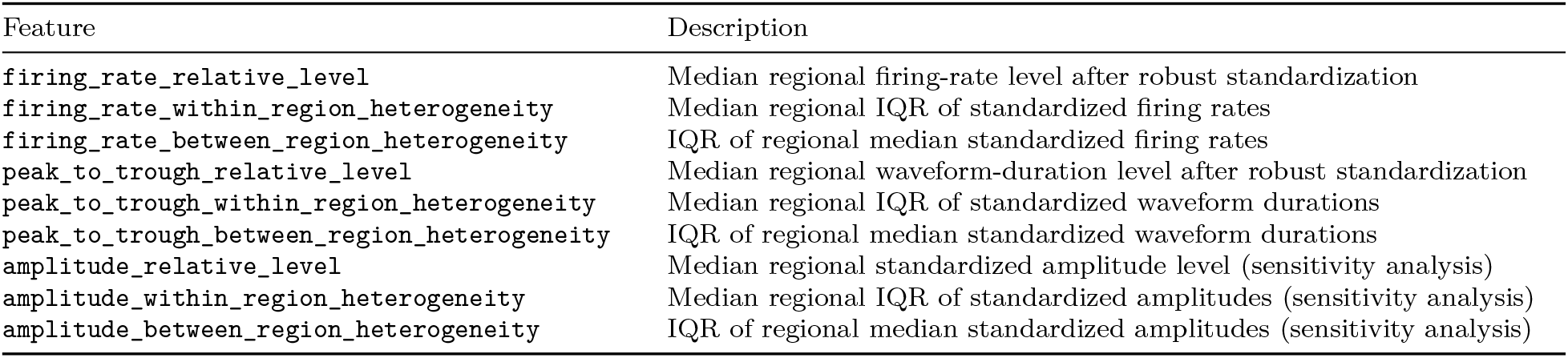
Neural features. The nine mouse-level neural features. The primary neural representationused the six firing-rate and waveform features; the three amplitude features were added in a sensitivity analysis. Session–region medians and IQRs were combined across sessions using medians before computing the mouse-level summaries.

